# Proximal immune-epithelial progenitor interactions drive chronic tissue sequelae post COVID-19

**DOI:** 10.1101/2023.09.13.557622

**Authors:** Harish Narasimhan, In Su Cheon, Wei Qian, Sheng’en Hu, Tanyalak Parimon, Chaofan Li, Nick Goplen, Yue Wu, Xiaoqin Wei, Young Min Son, Elizabeth Fink, Gislane Santos, Jinyi Tang, Changfu Yao, Lyndsey Muehling, Glenda Canderan, Alexandra Kadl, Abigail Cannon, Samuel Young, Riley Hannan, Grace Bingham, Mohammed Arish, Arka Sen Chaudhari, Jeffrey Sturek, Patcharin Pramoonjago, Yun Michael Shim, Judith Woodfolk, Chongzhi Zang, Peter Chen, Jie Sun

**Affiliations:** Beirne B. Carter Center for Immunology Research, University of Virginia, Charlottesville, VA 22908, USA; Division of Infectious Disease and International Health, Department of Medicine, University of Virginia, Charlottesville, VA 22908, USA; Department of Microbiology, Immunology and Cancer Biology, University of Virginia, Charlottesville, VA 22908, USA; Center for Public Health Genomics, University of Virginia School of Medicine, Charlottesville, VA 22908, USA; Women’s Guild Lung Institute, Department of Medicine, Cedars-Sinai Medical Center, Los Angeles, CA 90048, USA; Department of Biomedical Sciences, Cedars-Sinai Medical Center, Los Angeles CA 90048, USA; Robert and Arlene Kogod Center on Aging, Mayo Clinic, Rochester, MN 55905, USA; Department of Systems Biotechnology, Chung-Ang University, Anseong, Korea; Division of Asthma, Allergy and Immunology, Department of Medicine, University of Virginia, Charlottesville, VA 22908, USA; Division of Pulmonary and Critical Care Medicine, Department of medicine, University of Virginia, Charlottesville, VA 22908, USA; Department of Biomedical Engineering, University of Virginia, Charlottesville, VA 22908, USA; School of Medicine, University of Virginia, Charlottesville, VA 22908, USA

**Author notes:** **Co-correspondent authors: J.S.:****, P.C.:****, C.Z.:**. Co-first authors (Harish Narasimhan, In Su Cheon, Wei Qian, Sheng’en Hu, Tanyalak Parimon).

## Abstract

The long-term physiological consequences of SARS-CoV-2, termed Post-Acute Sequelae of COVID-19 (PASC), are rapidly evolving into a major public health concern. The underlying cellular and molecular etiology remain poorly defined but growing evidence links PASC to abnormal immune responses and/or poor organ recovery post-infection. Yet, the precise mechanisms driving non-resolving inflammation and impaired tissue repair in the context of PASC remain unclear. With insights from three independent clinical cohorts of PASC patients with abnormal lung function and/or viral infection-mediated pulmonary fibrosis, we established a clinically relevant mouse model of post-viral lung sequelae to investigate the pathophysiology of respiratory PASC. By employing a combination of spatial transcriptomics and imaging, we identified dysregulated proximal interactions between immune cells and epithelial progenitors unique to the fibroproliferation in respiratory PASC but not acute COVID-19 or idiopathic pulmonary fibrosis (IPF). Specifically, we found a central role for lung-resident CD8^+^ T cell-macrophage interactions in maintaining Krt8^hi^ transitional and ectopic Krt5^+^ basal cell progenitors, thus impairing alveolar regeneration and driving fibrotic sequelae after acute viral pneumonia. Mechanistically, CD8^+^ T cell derived IFN-γ and TNF stimulated lung macrophages to chronically release IL-1β, resulting in the abnormal accumulation of dysplastic epithelial progenitors and fibrosis. Notably, therapeutic neutralization of IFN-γ and TNF, or IL-1β after the resolution of acute infection resulted in markedly improved alveolar regeneration and restoration of pulmonary function. Together, our findings implicate a dysregulated immune-epithelial progenitor niche in driving respiratory PASC. Moreover, in contrast to other approaches requiring early intervention, we highlight therapeutic strategies to rescue fibrotic disease in the aftermath of respiratory viral infections, addressing the current unmet need in the clinical management of PASC and post-viral disease.

## INTRODUCTION

SARS-CoV-2 infection can lead to long-term pulmonary and extrapulmonary symptoms well beyond the resolution of acute disease, a condition collectively termed post-acute sequelae of COVID-19 (PASC) (1, 2). With effective treatment strategies and vaccines to tackle acute COVID-19, the emerging challenge is to manage chronic sequelae in the 60+ million people currently experiencing PASC (3, 4). Given the tropism of the virus to the respiratory tract, the lungs are particularly susceptible to extensive damage during primary infection, potentially resulting in chronic dyspnea, compromised lung function, and radiological abnormalities (1, 2). Notably, these sustained impairments of the lungs can persist up to 2 years post infection, unlike the majority of extrapulmonary sequelae which decline over time (4). Some individuals also develop a non-resolving fibroproliferative response – PASC pulmonary fibrosis (PASC-PF) and may require persistent oxygen supplementation and eventual lung transplantation (4–8). Despite progress in identifying dysregulated immune signatures associated with respiratory PASC (3, 7, 9–16), little is known regarding their mechanistic roles in driving pathological tissue remodeling, in part due to the lack of studies adopting “clinically relevant” animal models of post-viral lung disease.

It has been long established that type 2 alveolar epithelial (AT2) cells are the facultative stem cell of the lungs that undergo self-renewal and differentiate into type 1 alveolar (AT1) cells to replenish the denuded epithelial niche after alveolar injury (17–19). Recent work has further delineated the AT2 to AT1 trans-differentiation process and identified a transitional state characterized by high expression of cytokeratin 8 (Krt8^hi^), termed alveolar differentiation intermediates (ADI) (20), damage-associated transitional progenitors (DATP) (21), and pre-alveolar type-1 transitional cell state (PATS) (22). An analogous population of Krt17^+^Krt5^-^ aberrant basaloid cells has been described in idiopathic pulmonary fibrosis (IPF), which co-express several basal epithelial, mesenchymal and senescence markers (23, 24). Severe alveolar damage also induces the recruitment of Krt5^+^ basal cell progenitors to the distal lung, which persist in an undifferentiated state or differentiate into upper airway cell fates, causing alveolar “bronchiolization” (25–29). The accumulation of undifferentiated Krt8^hi^ transitional cells and/or the emergence of Krt5^-^Krt17^+^ aberrant basaloid cells and ectopic Krt5^+^ pods are hallmarks of lung injury, and their persistence have pathologically been associated with chronic diseases such as lung fibrosis (5, 30–37).

Currently, mechanisms underlying the long-term maintenance of these injury-associated dysplastic progenitors and their contributions to post-viral respiratory sequelae remain largely unknown. By comparing the pathological, immunological, and molecular features of various human respiratory PASC cohorts and mouse models of post-viral lung disease, we discovered spatially defined microenvironments characterized by dysregulated immune-epithelial progenitor interactions driving impaired alveolar regeneration and fibrotic outcomes after acute viral injury. Furthermore, we identify nodes for therapeutic intervention, which may be adopted in the clinic to mitigate chronic pulmonary sequelae after COVID-19.

## RESULTS

### Spatial association between dysplastic epithelial progenitors and CD8^+^ T cells is a hallmark of human PASC-PF

We examined diseased lung sections from two cohorts of PASC-PF patients that underwent lung transplantation at Cedars-Sinai Medical Center and University of Virginia. Patients had a mean age of 52.5 years and exhibited persistent pulmonary impairment and hypoxemia, requiring oxygen supplementation **(Supplementary Table 1)**. Lung histology was notable for extensive immune cell infiltration and collagen deposition in the alveolar epithelium **(Fig. 1a).** Given the heterogenous distribution of tissue pathology, we performed spatial transcriptomics on PASC-PF and control lungs to evaluate potentially distinct microenvironments between healthy and pathological areas of the lungs **(Extended data Fig. 1a)**. Using an agnostic evaluation of differentially regulated signaling pathways using Gene Set Enrichment Analysis (GSEA), we observed a robust upregulation of pathways associated with extracellular matrix deposition and fibrosis, as well as inflammation within PASC-PF lungs compared to controls **(Fig. 1b)**. Following UMAP visualization and clustering of the capture spots **(Extended data Fig. 1b,c)**, we observed that the expression of genes associated with AT1 and AT2 cells were reduced, whereas genes characterizing dysplastic epithelial cell states were elevated in PASC-PF lungs **(Fig. 1c, Extended data Fig. 1d)**. Consistent with the active inflammatory processes, CD8^+^ T cell gene expression was enriched in PASC-PF lungs **(Fig. 1c, Extended data Fig. 1d)**. Furthermore, we leveraged the power of spatial transcriptomics and found large areas of PASC-PF lungs populated by CD8^+^ T cells as well as areas of dysplastic repair whereas control lungs were primarily composed of alveolar epithelial cells **(Fig. 1d)**. Notably, the sporadic pockets of alveolar epithelium-rich areas in PASC-PF lungs were excluded from CD8^+^ T cells and dysplastic repair markers, which was particularly exemplified in the PASC-PF lung sample with a noticeably intact alveolar epithelium **(Fig. 1d)**. In accordance with this pattern, we observed a mild correlation between the distribution of CD8^+^ T cells and dysplastic epithelial progenitors within PASC-PF lungs **(Fig. 1e)**.

**Fig. 1.**
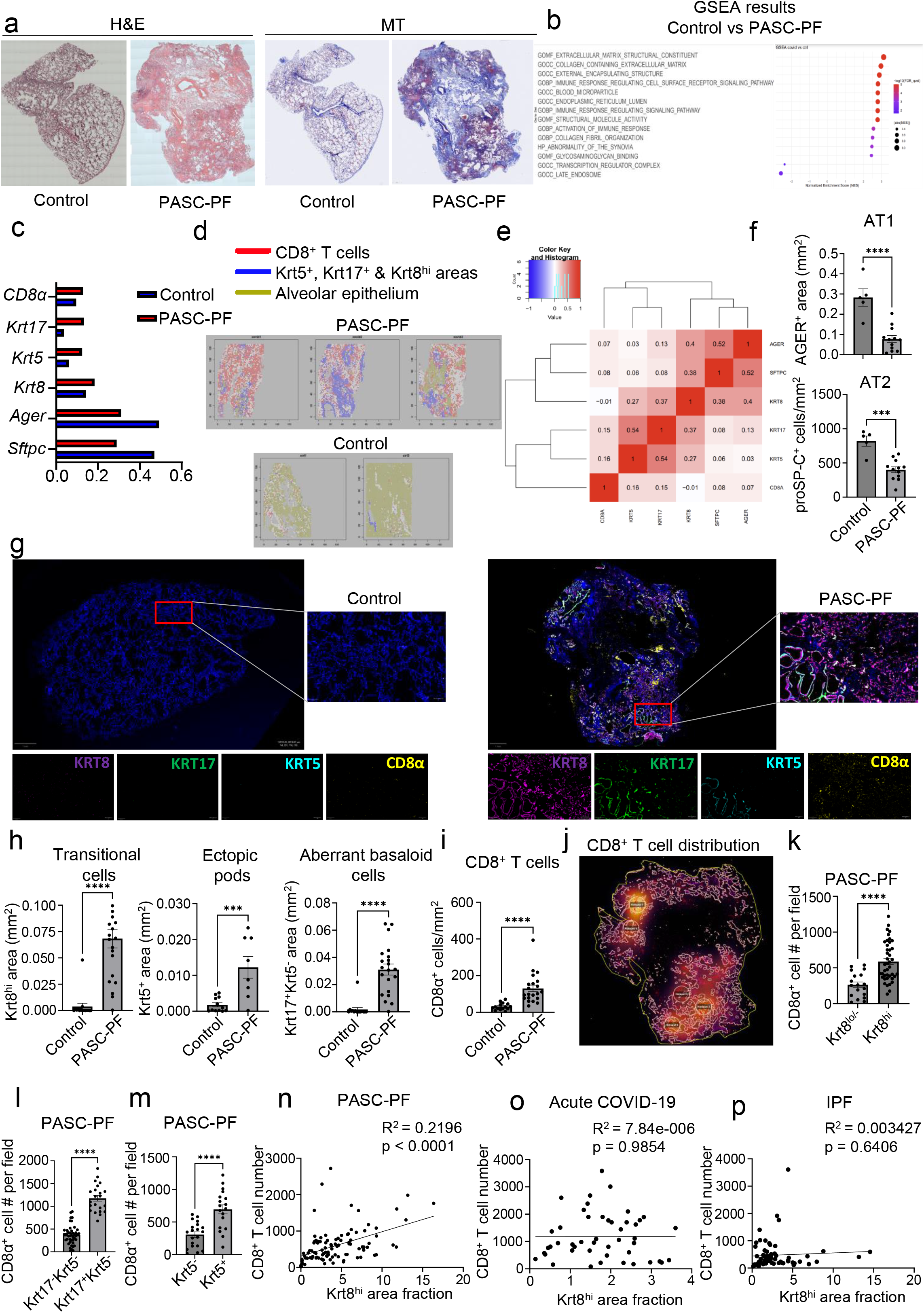
Spatial transcriptomics and imaging reveals chronic persistence and association between CD8^+^ T cells and epithelial progenitors in human PASC-PF. **a.** Representative hematoxylin & eosin (H&E) and Masson’s Trichrome (MT) images of control and PASC-PF lung sections. **b.** Unbiased GSEA analysis of signaling pathways differentially regulated between control and PASC-PF lungs. **c.** Quantification of the proportion of spots expressing genes characterizing various immune and epithelial populations in control and PASC-PF lungs. **d.** Visualization of CD8^+^ T cells, Krt5^+^, Krt8^hi^ and Krt17^+^-rich areas, and healthy alveolar epithelium based on gene expression signatures in human control and PASC-PF lungs. **e.** Heatmap of physical distribution of alveolar epithelium, dysplastic progenitors and CD8^+^ T cells. f. Quantification of AT1 (AGER^+^) and AT2 (proSP-C^+^) cells in control and PASC-PF lung sections (n = 5 control, 12 PASC-PF). **g.** Representative immunofluorescence images staining CD8^+^ T cells (CD8α^+^) and epithelial progenitors (Krt5^+^, Krt17^+^, and Krt8^hi^) in control and PASC-PF lung sections. **h.** Quantification of Krt8^hi^, Krt5^+^, Krt17^+^Krt5^-^ and **i.** CD8^+^ T cells in control and PASC-PF lung sections (n = 15 control, 21 PASC-PF). **j.** Unbiased analysis of CD8^+^ T cell distribution in a PASC-PF lung section using QuPath. **k.** Quantification of the distribution of CD8^+^ T cells between fields with or without Krt8^hi^, **l.** Krt5^-^Krt17^+^, and **m.** Krt5^+^ areas in human PASC-PF lungs. **n.** Simple linear regression of CD8^+^ T cell number and Krt8^hi^ area fraction in human PASC-PF, **o.** acute COVID-19, and **p.** IPF lung sections. Data are expressed as mean ± SEM. Statistical analyses were conducted using a two-tailed unpaired t-test (e,g,h,j-m). *p < 0.05; **p < 0.01; ***p < 0.001; ****p < 0.0001.

Although the spatial transcriptomics findings suggested potential immune-epithelial progenitor crosstalk, they may be confounded by the limited capture area, lack of single-cell resolution as well as the spectrum of gene expression observed across epithelial progenitor states (36). Moreover, only one of three PASC-PF sampled sections contained prominent alveolar epithelium-rich areas, whereas the controls lacked any significant representation of dysplastic epithelial progenitors **(Fig. 1d).** These limitations hindered further analyses on the spatial distribution of immune cells within healthy and dysplastic areas. Thus, to address these caveats, we validated our findings in a larger number of PASC-PF and control lungs via multicolor immunofluorescence.

Consistent with the spatial transcriptomics data, we found reduced levels of AT1 and AT2 cell markers in PASC-PF lungs compared to controls, indicating a persistent defect in alveolar regeneration **(Fig. 1f, Extended data Fig. 2a)** (38). We also observed chronic persistence of ectopic Krt5^+^ pods, Krt8^hi^ transitional cells, and Krt5^-^Krt17^+^ aberrant basaloid cells in PASC-PF lungs compared to controls, which is concordant with recent reports (5, 8, 32, 39) **(Fig. 1g,h)**. Moreover, we found that high KRT8 expression was inclusive of all dysplastic epithelial progenitors including transitional cells, Krt17^+^Krt5^-^ aberrant basaloid and Krt5^+^ metaplastic basal cells **(Fig. 1g, Extended data Fig. 2b)** (35). PASC-PF lungs harbored widespread expression of alpha smooth muscle actin (αSMA), indicative of myofibroblast activity, in proximity to areas of dysplastic repair (**Extended data Fig. 2c**) (23, 24, 36). Therefore, PASC-PF is characterized by the sustained loss of functional alveolar epithelial cells, and the persistence of metaplastic Krt5^+^ cells, Krt8^hi^ transitional cells, and Krt5^-^Krt17^+^ aberrant basaloid cells, which is histologically akin to other fibrotic lung diseases such as idiopathic pulmonary fibrosis (IPF) **(Extended data Fig. 2b,c)** (30, 31).

Previously, we reported that increased CD8^+^ T cell levels in the bronchoalveolar lavage (BAL) fluid were associated with impaired lung function in COVID-19 convalescents (9, 10). As observed with spatial transcriptomics, CD8^+^ T cell numbers were elevated in PASC-PF patient lungs **(Fig. 1g,i)** as well as in acute COVID-19, but not in IPF lungs when compared to controls **(Extended data Fig. 2d,e)**. The mild correlation in localization revealed by spatial transcriptomics was corroborated by a striking spatial association between CD8^+^ T cells, and Krt8^hi^, Krt17^+^Krt5^-^, and Krt5^+^ areas representing dysplastic repair upon analysis of their distribution in PASC-PF lungs by microscopy **(Fig. 1j-m, Extended data Fig. 3a-c)**. Notably, this correlation between CD8^+^ T cells, and dysplastic epithelial progenitors was unique to PASC-PF lungs but not seen in lungs from control, acute COVID-19 or IPF conditions **(Fig. 1n-p, Extended data Fig. 3d-k)**. Collectively, these data indicate that the spatiotemporal colocalization of CD8^+^ T cells and areas of dysplastic repair is a unique feature of post-viral pulmonary fibrosis and supports immune-epithelial progenitor interactions potentially contributing to the observed defects in alveolar regeneration and chronic pulmonary sequelae.

### A mouse model of post-viral lung sequelae recapitulating features of human PASC-PF

To investigate the role of immune-epithelial progenitor interactions, we aimed to develop a mouse model to capture the cellular and pathological features observed in PASC-PF lungs. We used a mouse-adapted (MA-10) strain of the SARS-CoV-2 virus to productively infect WT mice. Notably, SARS-CoV-2 MA-10 infection is known to induce acute lung disease and pneumonia in mice, characterized by substantial damage to the airway epithelium, fibrin deposition, and pulmonary edema (40). Since aging is associated with an increased propensity to develop lung fibrosis post viral injury as well as severe disease after SARS-CoV-2 infection in mice, we included both young and aged mice in our study (41–43). As expected, aged C57BL/6 mice infected with SARS-CoV-2 MA-10 had increased morbidity and mortality compared to young mice **(Fig. 2a, Extended data Fig. 4a)**. Indeed, we observed marked inflammation and tissue damage acutely (at 10 days post infection (dpi)) (**Extended data Fig. 4b)** (40, 44). However, irrespective of age, the majority of the lungs recovered from the acute damage and only moderate pathology restricted to the subpleural regions was observed at the chronic phase (35dpi) of infection **(Fig. 2b, Extended data Fig. 4c)**.

**Fig. 2.**
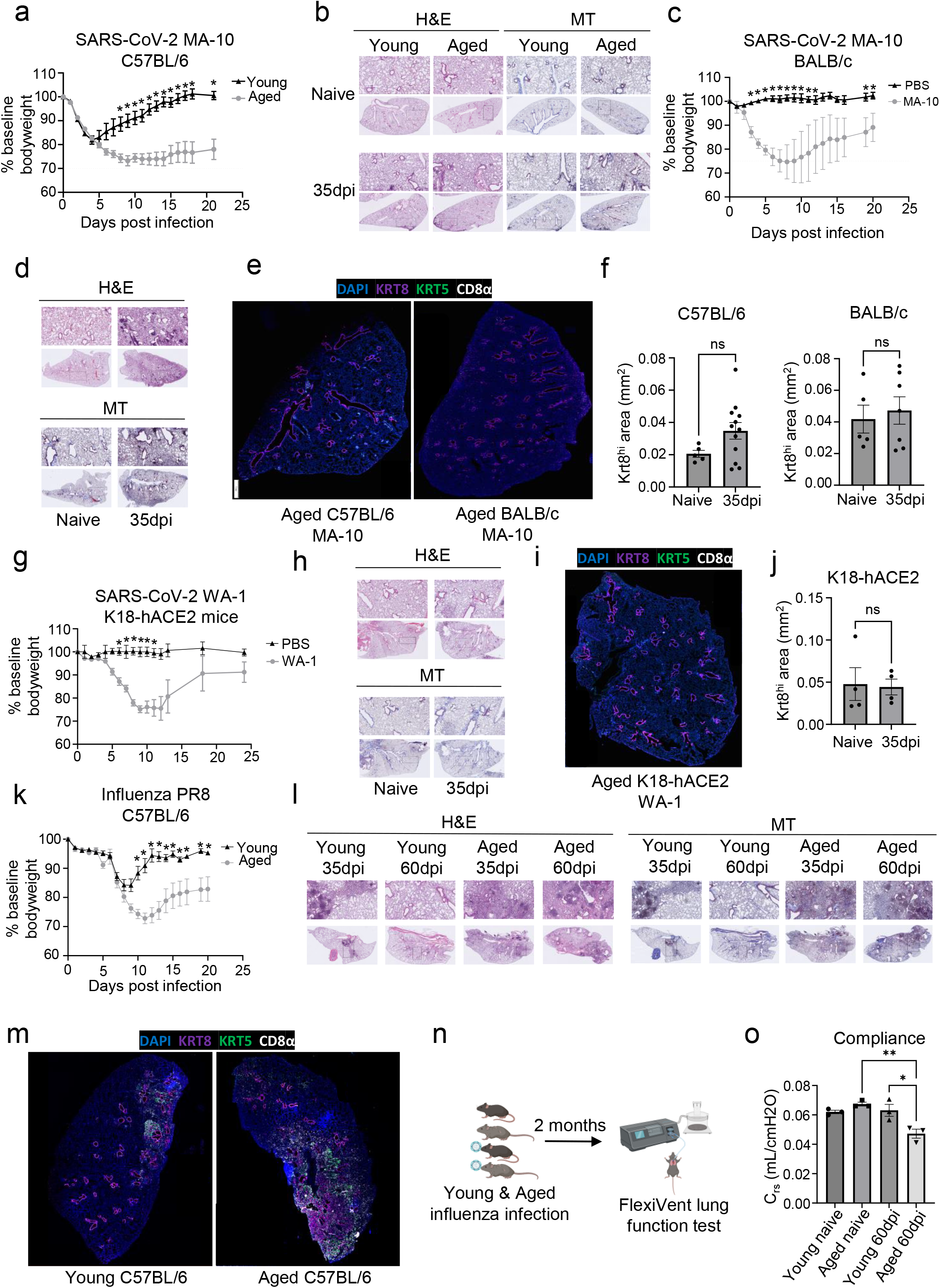
Development of a mouse model of post-viral fibrosis recapitulating features of human PASC-PF. **a.** Percent change in bodyweight of young (8-10 weeks) and aged (20-22 months) C57BL/6 mice post infection with SARS-CoV-2 MA-10 virus. **b.** Representative H&E and MT images of young and aged C57BL/6 lungs post SARS-CoV-2 infection (35dpi) or PBS administration. **c.** Percent change in bodyweight of aged (12-14 months) BALB/c mice post infection with SARS-CoV-2 MA-10 virus or PBS administration. **d.** Representative H&E and MT images of aged BALB/c lungs post SARS-CoV-2 infection (35dpi) or PBS administration. **e.** Representative immunofluorescence images staining CD8^+^ T cells (CD8α) and epithelial progenitors (Krt5^+^ and Krt8^hi^) in aged C57BL/6 and BALB/c mouse lungs post SARS-CoV-2 infection (35dpi). **f.** Quantification of Krt8^hi^ area in aged C57BL/6 and BALB/c mice post SARS-CoV-2 infection (35dpi). **g.** Percent change in bodyweight of aged (12-14 months) K18-hACE2 mice post infection with SARS-CoV-2 WA-1 virus or PBS administration. **h.** Representative H&E and MT images of aged K18-hACE2 lungs post SARS-CoV-2 infection (35dpi) or PBS administration. **i.** Representative immunofluorescence image staining CD8^+^ T cells (CD8α) and epithelial progenitors (Krt5^+^ and Krt8^hi^) in aged K18-hACE2 mouse lungs post SARS-CoV-2 infection (35dpi). **j.** Quantification of Krt8^hi^ area in aged K18-hACE2 mice post SARS-CoV-2 infection (35dpi). **k.** Percent change in bodyweight of young (8-10 weeks) and aged (20-22 months) C57BL/6 mice post infection with 75PFU of PR8 influenza virus. **l.** Representative H&E and MT images of young and aged C57BL/6 lungs post PR8 influenza virus infection (35dpi and 60dpi). **m.** Representative immunofluorescence images staining CD8^+^ T cells (CD8α) and epithelial progenitors (Krt5^+^ and Krt8^hi^) in young and aged C57BL/6 mouse lungs post H1N1 influenza A/PR8/34 virus infection (35dpi). **n.** Schematic of the FlexiVent system to evaluate pulmonary function in mice. **o.** Evaluation of compliance of the respiratory system in young and aged mice post PBS administration or influenza infection (60dpi). Data are expressed as mean ± SEM. Statistical analyses were conducted using a two-tailed unpaired t-test (f,j), multiple *t* tests (a,c,g,k), and an ordinary one-way ANOVA (o). *p < 0.05; **p < 0.01.

Next, we tested SARS-CoV-2 MA-10 infection in aged BALB/c mice, which was previously reported to induce more robust inflammation and fibrotic sequelae compared to C57BL/6 mice (41) **(Fig. 2c, Extended data Fig. 4e)**. Consistent with the report, we observed persistent immune cell infiltration, which was largely restricted to the peri-bronchiolar regions at 35dpi **(Fig. 2d)**. However, similar to the aged C57BL/6 mice, minimal signs of collagen deposition were observed at later time points in the alveolar epithelium **(Fig. 2d, Extended data Fig. 4g)**. Consistent with what was previously reported (41), BALB/c mice failed to maintain pulmonary CD8^+^ T cells **(Extended data Fig. 4d,h)**, a prominent feature of human PASC-PF(**Fig. 1d)**. Moreover, no significant difference was observed in the development and persistence of Krt8^hi^ transitional cells between aged naïve and infected (35dpi) mice in both genetic backgrounds in spite of substantial alveolar damage during acute disease **(Fig. 2e,f, Extended data Fig. 4i-k)**.

Finally, we infected transgenic mice expressing the human ACE2 receptor under the cytokeratin 18 promoter (K18-hACE2) with the WT SARS-CoV-2 virus (USA-WA1/2020), which is known to result in perivascular and alveolar inflammation during acute disease (45). Although infection induced significant morbidity and mortality following infection, minimal signs of alveolar injury were observed at chronic timepoints **(Fig. 2g,h, Extended data Fig. 4l,m)**. Consistent with the histopathological findings, no appreciable difference was observed in CD8^+^ T or Krt8^hi^ transitional cells between infected and naïve mice **(Fig 2i,j, Extended data Fig. 4n)**. Thus, SARS-CoV-2 infection in these three mouse models failed to recapitulate key features of tissue pathology and dysplastic lung repair observed in human PASC-PF.

Previously, we and others reported persistent lung inflammation and tissue pathology after influenza viral infection, particularly in aged C57BL/6 mice (9, 42, 46). Therefore, we infected young and aged WT C57BL/6 mice with influenza H1N1 A/PR8/34 strain, which causes substantial viral pneumonia and alveolar damage during acute disease (42, 47). Similar to SARS-CoV-2 infection, aged mice exhibited increased morbidity and mortality post influenza infection compared to young mice **(Fig. 2k, Extended data Fig. 4o)**. In contrast to SARS-CoV-2 infection, we observed persistent immune cell infiltration and collagen deposition in the alveolar epithelium, particularly in aged mice that persisted to 60dpi **(Fig. 2l, Extended data Fig. 4p)**. Moreover, lungs from aged mice harbored significantly larger Krt5^+^ and Krt8^hi^ areas of dysplastic repair as well as higher levels of CD8^+^ T cells post influenza viral pneumonia, similar to human PASC-PF lungs **(Fig. 2m, 3b-d).** Although the size of dysplastic repair areas diminished over time, substantial immune cell infiltration and collagen deposition was observed up to 8 months post infection **(Extended data Fig. 5a,b)**. The alveolar architecture was disrupted within these areas of fibroproliferation, which harbored high levels of CD8^+^ T cells and dysplastic epithelial progenitors, indicating prolonged inflammation and pathological remodeling persist up to 250dpi **(Extended data Fig. 5b-f)** (48). The observed cellular changes were associated with sustained defects in pulmonary function in aged mice up to 60dpi following influenza viral pneumonia, mimicking the sustained impairment in lung function observed in respiratory PASC patients **(Fig. 2n,o)** (9, 39). Although the exact mechanisms underlying the divergent trajectories in recovery following acute alveolar injury due to SARS-CoV-2 MA-10 and influenza PR8 infections remain unclear, extensive evaluation of various mouse viral pneumonia models indicated that influenza infection in aged mice has closer histopathological alignment with features of persistent pulmonary sequelae observed in PASC-PF lungs. Thus, influenza infection of aged mice can serve as a clinically relevant model to study the mechanisms of viral infection-mediated lung fibrosis.

### Exuberant tissue CD8^+^ T cell responses impair alveolar regeneration and promote dysplastic lung repair following viral pneumonia

To further investigate the spatiotemporal dynamics of lung fibrosis, we infected young and aged mice with influenza virus and characterized the immune and epithelial progenitor compartments over time **(Fig 3a, Extended data Fig. 6a)**. The induction of epithelial progenitor activity was comparable in young and aged mice during the acute phase of infection (up to 14dpi) but diverged at later timepoints with increased Krt5^+^ and Krt8^hi^ cells in aged mice **(Fig. 3a-c)**. These trends correlated with a persistent age-associated defect in alveolar regeneration, exemplified by a sustained reduction in AT2 cell numbers **(Extended data Fig. 6b-d)**. Consistent with our previous study, lungs from aged mice harbored significantly higher levels of CD8^+^ T cells post influenza infection compared those of young mice **(Fig. 3d)** (9, 42). Similar to human PASC-PF lungs, we also observed a spatial association between CD8^+^ T cells and Krt8^hi^ areas of dysplastic repair, reinforcing the relevance of this model to study post-viral pulmonary sequelae (**Fig. 3e, 1j**). Furthermore, the association between CD8^+^ T cells and areas of dysplastic repair was seen only at post-acute timepoints and strengthened over time, recapitulating features of human lungs after severe SARS-CoV-2 infection and suggesting these immune-epithelial progenitor interactions are primarily a feature of chronic sequelae of viral infections (**Fig. 3f, Extended data Fig. 6e-g).**

**Fig. 3.**
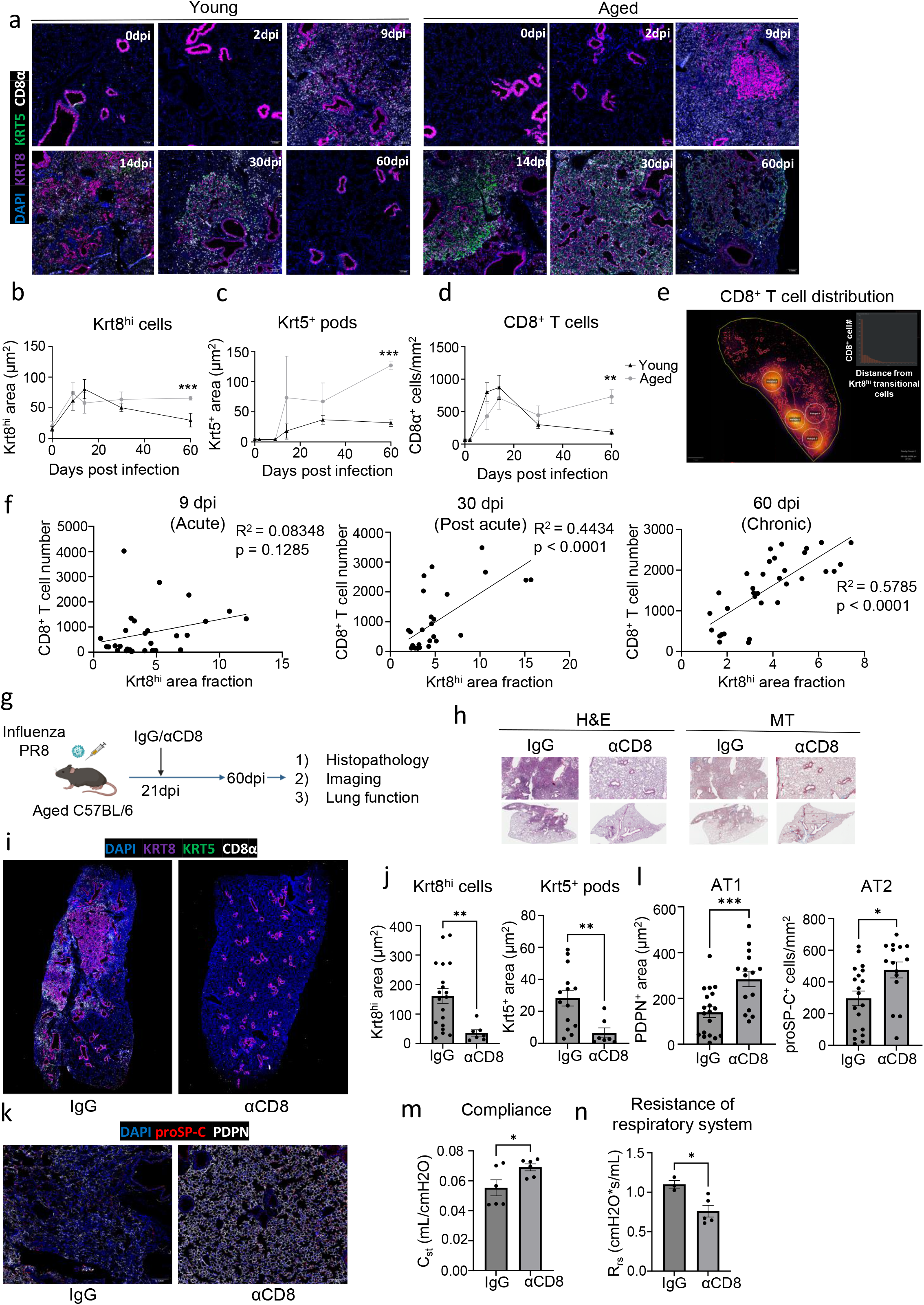
Persistent CD8^+^ T cell activity impairs functional alveolar regeneration and promote dysplastic lung repair. **a.** Representative immunofluorescence images staining CD8^+^ T cells (CD8α^+^) and epithelial progenitors (Krt5^+^ and Krt8^hi^) in young and aged C57BL/6 mouse lungs over the course of influenza infection. **b.** Quantification of Krt8^hi^, **c.** Krt5^+^, and **d.** CD8^+^ T cells in young and aged lungs post influenza infection (n = 4 per time point). **e.** Unbiased analysis of CD8^+^ T cell distribution in an aged mouse lung section post influenza infection (60dpi). **f.** Simple linear regression of CD8^+^ T cell number and Krt8^hi^ area fraction in influenza-infected mice at 9, 30, and 60dpi. **g.** Experimental design for CD8^+^ T cell depletion post influenza infection. **h.** Representative H&E and MT images of aged C57BL/6 lungs post influenza infection (60dpi) treated with αCD8 or control IgG antibody (Ab). **i.** Representative immunofluorescence images staining CD8^+^ T cells (CD8α^+^) and epithelial progenitors (Krt5^+^ and Krt8^hi^) in aged C57BL/6 mouse lungs post influenza infection (60dpi) treated with αCD8 or control IgG Ab. **j.** Quantification of Krt8^hi^ and Krt5^+^ area in aged influenza-infected lung sections treated with αCD8 or control IgG Ab. (n = 19 control IgG, 7 αCD8). **k.** Representative immunofluorescence images staining AT1 (PDPN^+^) and AT2 (proSP-C^+^) in aged C57BL/6 mouse lungs post influenza infection (60dpi), treated with αCD8 or control IgG Ab. **l.** Quantification of AT1 (PDPN^+^) and AT2 (proSP-C^+^) cells in aged influenza-infected lung sections treated with αCD8 or control IgG Ab. (n = 19 control IgG, 15 αCD8). **m.** Evaluation of static compliance (C_st_) and **n.** resistance of the respiratory system (R_rs_) in aged C57BL/6 mouse lungs post influenza infection (60dpi), treated with αCD8 or control IgG Ab. Data are expressed as mean ± SEM. Statistical analyses were conducted using a two-tailed unpaired t-test (j,l,m,n) and multiple *t* tests (b-d). *p < 0.05; **p < 0.01; ***p < 0.001.

To understand the role of the persistent CD8^+^ T cells at this stage, we treated aged influenza-infected mice with CD8^+^ T cell-depleting Ab (αCD8) or isotype control Ab starting from 21dpi **(Fig. 3g)**. This post-acute timepoint was chosen to ensure no interference with the essential antiviral activities of CD8^+^ T cells during acute infection as the virus is completely cleared by 15dpi in this model (42). CD8^+^ T cell depletion improved histological evidence of disease in aged mice **(Fig. 3h, Extended data Fig. 7a).** Importantly, Krt5^+^ and Krt8^hi^ areas were significantly reduced by depletion of CD8^+^ T cells **(Fig. 3i,j),** suggesting that CD8^+^ T cells are essential for the maintenance of dysplastic epithelial progenitors after recovery from acute disease. Interestingly, AT1 and AT2 cells were markedly increased, and the alveolar architecture was restored after CD8^+^ T cell depletion **(Fig. 3k,l)**. We also observed a concomitant improvement in lung function after αCD8 treatment, suggesting that exuberant CD8^+^ T cell responses in the aftermath of acute disease affected the restoration of alveolar spaces, resulting in chronic impairment of alveolar gas-exchange function **(Fig. 3m,n).** Notably, depletion of CD8^+^ T cells resolved Krt5^+^ but not Krt8^hi^ areas in young mice and did not dramatically affect lung pathology or alveolar regeneration (**Extended data Fig. 7b-g)**, suggesting that CD8^+^ T cells may specifically influence age-associated dysplastic lung repair.

To dissect the roles of circulating CD8^+^ T cells and lung resident memory CD8^+^ T cells in impairing alveolar regeneration, we used low and high dose αCD8 treatment to deplete circulating and pulmonary CD8^+^ T cells respectively, in aged influenza-infected mice (42, 49). We found that the resolution of areas of dysplastic repair only occurs upon depletion of pulmonary CD8^+^ T cells but not circulating CD8^+^ T cells **(Extended data Fig. 7h,i)** (42), suggesting that lung-resident CD8^+^ T cells are required for the maintenance of dysplastic epithelial progenitors. Taken together, our results strongly implicate the activity of CD8^+^ T cells persisting in the lungs in the aftermath of viral pneumonia in the development of chronic pulmonary sequelae.

### Spatial transcriptomics reveal proximal interactions between CD8^+^ T cells, macrophages, and Krt5^+^ and Krt8^hi^-rich areas of dysplastic repair

To capture the spatially confined proximal interactions between immune and epithelial progenitor cells within areas of dysplastic repair, we performed spatial transcriptomics on aged influenza-infected mouse lungs (60dpi) treated with control Ab or αCD8 **(Fig. 4a)**. Following UMAP visualization and clustering of the capture spots **(Extended data Fig. 8a,b)**, we observed a strong association between gene expression signatures of CD8^+^ T cells, and Krt5^+^ and Krt8^hi^ areas of dysplastic repair, similar to immunostaining results and human PASC-PF lungs **(Fig 4b, 3e)**. Consistent with previous data, gene expression signatures of Krt5^+^ and Krt8^hi^ progenitors were dramatically reduced in αCD8-treated lungs, with a concomitant increase in healthy alveolar epithelial cells **(Fig 4b, 3i-k, Extended data Fig. 8d,e)**. These findings also corresponded with the expression pattern of a gene-set indicative of fibrosis, with the strongest enrichment within Krt5^+^ and Krt8^hi^ areas and a similar reduction in αCD8-treated lungs **(Extended data Fig. 8c)**. We further performed an agnostic evaluation of signaling pathways differentially regulated within the healthy alveolar epithelium (AE) compared to Krt5^+^ and Krt8^hi^-rich areas of dysplastic repair (Krt) **(Fig. 4c)**. Several pathways associated with inflammatory responses were highly active within areas of dysplastic repair, whereas growth factor responses and restoration of the vasculature were prominently observed in the healthy alveolar epithelium.

**Fig. 4.**
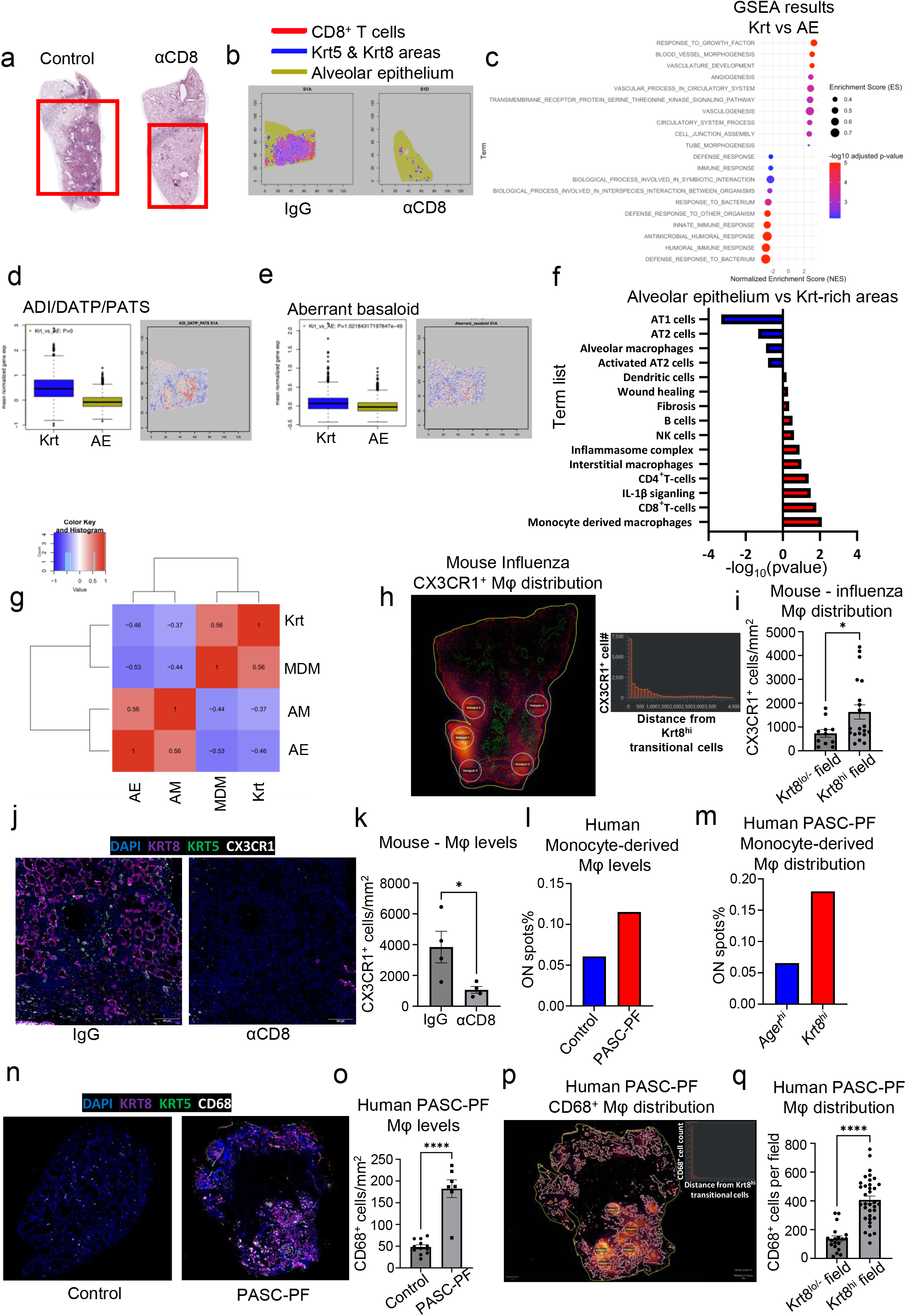
Spatial transcriptomics reveal a CD8^+^ T cell-macrophage-epithelial progenitor niche that drives dysplastic lung repair. **a.** Representative H&E images of aged influenza-infected mice (60dpi) treated with control IgG Ab or αCD8 that were mounted on the 10X Visium slide. **b.** Visualization of CD8^+^ T cells, Krt5^+^ and Krt8^hi^ transitional cells, and alveolar epithelial cells based on gene signatures. **c.** Unbiased GSEA analysis of signaling pathways enriched in Krt5^+^ and Krt8^hi^-rich areas compared to healthy alveolar epithelium. **d.** Boxplots comparing gene expression signature of ADI/DATP/PATS and **e.** aberrant basaloid cells in Krt5^+^ and Krt8^hi^-rich areas and healthy alveolar epithelium. **f.** Gene expression signatures of various immune and epithelial cell types as well as signaling pathways in Krt5^+^ and Krt8^hi^-rich areas and healthy alveolar epithelium. **g.** Heatmap of the physical distribution of monocyte-derived macrophages (MDM), alveolar macrophages (AM), Krt5^+^ and Krt8^hi^ areas (Krt), and healthy alveolar epithelium (AE) in aged influenza-infected (60dpi) mouse lungs. **h.** Unbiased analysis of the distribution of CX3CR1^+^ macrophages in an aged influenza-infected (60dpi) mouse lung section. **i.** Quantification of CX3CR1^+^ cells within Krt8^-/lo^ and Krt8^hi^ fields in aged influenza-infected (60dpi) lungs (n=5). **j.** Representative immunofluorescence images of aged influenza-infected (60dpi) lungs treated with control IgG Ab or αCD8, staining monocyte-derived macrophages (CX3CR1) and epithelial progenitors (Krt5 and Krt8). **k.** Quantification of CX3CR1^+^ macrophages in aged influenza-infected (60dpi) lungs treated with control IgG Ab or αCD8 (n=4). **l.** Quantification of the proportion of spots expressing the gene signature of monocyte-derived macrophages in human control and PASC-PF lungs. **m.** Quantification of the proportion of spots expressing the gene signature of monocyte-derived macrophages in *Ager^hi^* and *Krt8^hi^*areas within human PASC-PF lungs. **n.** Representative immunofluorescence images of human PASC-PF lungs staining macrophages (CD68) and epithelial progenitors (Krt5 and Krt8). **o.** Quantification of CD68^+^ macrophages in human control and PASC-PF lungs (n=11 control; 7 PASC-PF). **p.** Unbiased analysis of the distribution of CD68^+^ macrophages in a human PASC-PF lung using QuPath. **q.** Quantification of CD68^+^ macrophages within Krt8^-/lo^ and Krt8^hi^ fields in human PASC-PF lungs (n=6). Data are expressed as mean ± SEM. Statistical analyses were conducted using a two-tailed unpaired t-test (k,o,q). *p < 0.05; **p < 0.01.

Upon investigation of specific gene expression-signatures, we observed an enrichment of the ADI/DATP/PATS (20–22) signature characterizing Krt8^hi^ transitional cells **(Fig. 4d)** as well as the analogous population of aberrant basaloid cells observed in humans (23, 24) **(Fig. 4e)** particularly within areas of dysplastic repair. Evaluation of various immune and epithelial cell marker signatures revealed prominent enrichment of monocyte-derived macrophages in Krt5^+^ and Krt8^hi^-rich areas in addition to other immune cells including CD4^+^ T cells, interstitial macrophages, B-cells, and natural killer cells **(Fig. 4f, Extended data Fig. 8f,g)**. In contrast, gene expression signatures associated with pro-repair tissue-resident alveolar macrophages, AT1, and AT2 cells were primarily observed within the healthy alveolar epithelium **(Fig 4f, Extended data Fig. 8e,h)**. To further characterize the immune-epithelial progenitor niche, we performed a correlation analysis and found that CD8^+^ T cells and monocyte-derived macrophages were physically clustered around Krt5^+^ and Krt8^hi^ areas of dysplastic repair and excluded from areas enriched with alveolar macrophages and mature alveolar epithelial cells **(Fig. 4g)**.

To validate our findings from the spatial transcriptomics data, we immunostained aged influenza-infected mouse lungs and identified a similar enrichment of CX3CR1^+^ monocyte-derived macrophages within Krt8^hi^ areas of dysplastic repair **(Fig. 4h,i)**. Interestingly, CD8^+^ T cell depletion resulted in a decrease in CX3CR1^+^ macrophage numbers, suggesting a potential role for their recruitment and maintenance in lungs **(Fig. 4j,k, Extended data Fig. 8g)**. Upon further analysis of the human spatial transcriptomics data, we observed a similar increase in the gene signature of monocyte-derived macrophages within PASC-PF lungs **(Fig. 4l)**, which were particularly localized within Krt8^hi^ areas of dysplastic repair **(Fig. 4m, Extended data Fig. 9a-c)**. We further validated this by immunostaining, where human PASC-PF lungs exhibited an increase in macrophage populations compared to controls **(Fig. 4n,o)**, which were also strongly enriched within areas of dysplastic repair **(Fig. 4p,q)**. Together, our data revealed a conserved finding in both mouse and human post-viral lungs, where CD8^+^ T cells are present in fibrotic regions and in proximity to fibroproliferative mediators such as monocyte-derived macrophages, and dysplastic epithelial progenitors, representing a pathological niche after acute respiratory viral infections.

### A CD8^+^ T cell-macrophage axis induces IL-1β release to arrest AT2 trans-differentiation in the transitional cell state

We postulated the interactions between CD8^+^ T cells, macrophages, and epithelial progenitors within this pathological niche generated molecular cues to create a profibrotic microenvironment. We observed an enrichment of IL-1R signaling as well as inflammasome signatures in the areas of dysplastic repair compared to the healthy alveolar epithelium in human PASC-PF **(Fig 5a, Extended data Fig. 10a)**, which was conserved with aged influenza-infected mouse lungs as well **(Fig. 5b, Extended data Fig. 10b)**.

**Fig. 5.**
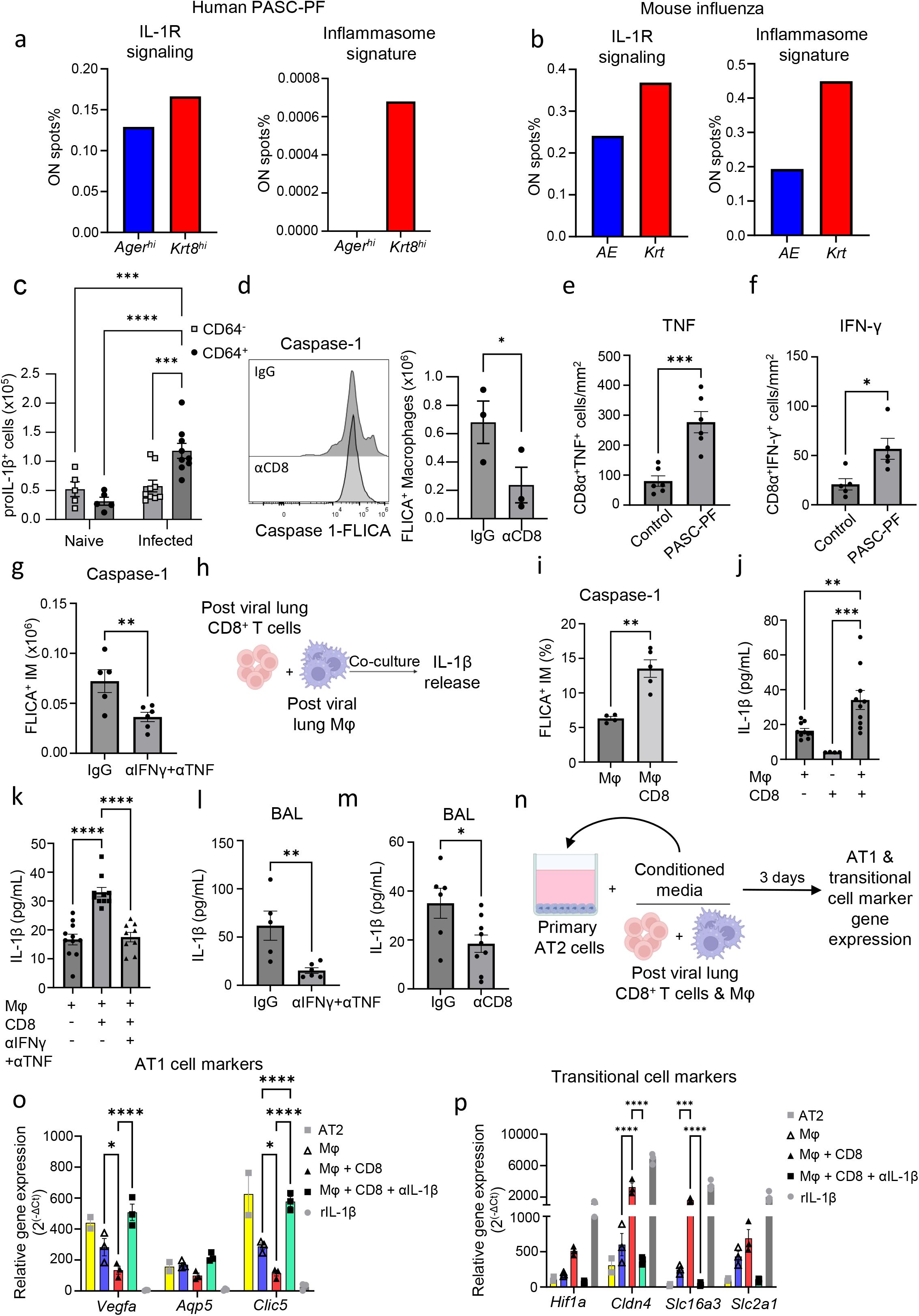
CD8+ T cells promote macrophage-mediated IL-1β release via IFN-γ and TNF. **a.** Quantification of the proportion of spots expressing the gene signature characterizing IL-1R signaling and inflammasome activity in *Ager^hi^* and *Krt8^hi^* areas within human PASC-PF lungs. **b.** Quantification of the proportion of spots expressing the gene signature characterizing IL-1R signaling and inflammasome activity within healthy (*AE*) and dysplastic (*Krt*) areas of aged influenza-infected (60dpi) mouse lungs. **c.** Quantification of proIL-1β^+^ cells in aged influenza-infected mice (42dpi) via flow cytometry. **d.** Assessment of caspase-1 activity in the lungs of aged influenza-infected mice (42dpi) after treatment with control IgG Ab or αCD8 by FLICA. **e.** Quantification of TNF^+^ and **f.** IFN-γ^+^ CD8^+^ T cells in human control and PASC-PF lungs. **g.** Assessment of caspase-1 activity in the lungs of aged influenza-infected mice (42dpi) after treatment with control IgG Ab or αIFN-γ+αTNF by FLICA. **h.** Schematic of the *ex vivo* macrophage and CD8^+^ T cell coculture system. **i.** Assessment of caspase-1 activity following coculture by FLICA. **j.** Evaluation of IL-1β release into supernatant following coculture by ELISA. **k.** Evaluation of IL-1β release into supernatant following *in vitro* IFN-γ and TNF blockade in the coculture system. **l.** Evaluation of IL-1β levels in BAL fluid of aged influenza-infected mice (42dpi) after treatment with control IgG Ab or neutralizing αIFN-γ+αTNF. **m.** Evaluation of IL-1β levels in BAL fluid of aged influenza-infected mice (42dpi) after treatment with control IgG Ab or αCD8. **n.** Experimental design for 2D culture of primary murine AT2 cells for 3 days without/with conditioned media from macrophages, macrophages + CD8^+^ T cells, macrophages + CD8^+^ T cells + αIL-1β, and rIL-1β (positive control). **o.** Gene expression of AT1 cell markers 3 days post AT2 cell culture following exposure to conditioned media. **p.** Gene expression of transitional cell markers 3 days post AT2 cell culture following exposure to conditioned media. Data are expressed as mean ± SEM. Statistical analyses were conducted using a two-tailed unpaired t-test (d-g, i-m), a two-way ANOVA (c), and a one-way ANOVA (o,p). *p < 0.05; **p < 0.01; *** p < 0.001; ****p < 0.0001.

IL-1β has been shown to promote the expansion of transitional Krt8^+^ cells upon bleomycin injury (21, 50). Therefore, we investigated if IL-1β mediated the development of post-viral pulmonary fibrosis. First, we found that CD64^+^ macrophages were major producers of pro-IL-1β compared to CD64^-^ cells in influenza-infected aged lungs **(Fig. 5c, Extended data Fig. 10c)**. Since mature IL-1β release from cells requires caspase-1 mediated pro-IL-1β cleavage (51), we examined caspase-1 activity in lung macrophages after CD8^+^ T cell depletion. Strikingly, CD8^+^ T cell depletion significantly reduced caspase-1 activity in lung macrophages **(Fig. 5d).** Moreover, inflammasome gene signatures were attenuated with αCD8 treatment (**Extended data Fig. 10d)**, supporting the notion that CD8^+^ T cells persisting in the lungs post infection promote chronic inflammasome activation and IL-1β release by macrophages.

CD8^+^ T effector and memory T cells are known to express high levels of IFN-γ and TNF (52). Indeed, PASC-PF lungs were found to harbor high levels of IFN-γ^+^ and TNF^+^ CD8^+^ T cells, similar to aged influenza-infected lungs **(Fig. 5e,f, Extended data Fig.11 a-c)**(3). Moreover, areas of dysplastic repair in particular were enriched with IFN-γ and TNF signaling in human PASC-PF lungs **(Extended data Fig. 11d)**. To determine whether IFN-γ and TNF mediate the observed inflammasome activation, we treated aged-influenza infected mice with neutralizing antibodies against both IFN-γ and TNF starting 21dpi and observed a decrease in caspase-1 activity **(Fig. 5g)**. As this effect was similar to αCD8-treatment **(Fig. 5d, Extended data Fig. 10d)**, we directly tested if CD8^+^ T cell-derived IFN-γ and TNF regulated macrophage inflammation activation. We used an *ex vivo* coculture of macrophages and CD8^+^ T cells isolated from mice previously infected with influenza (42dpi) to assess IL-1β release **(Fig. 5h)**. Indeed, we observed CD8^+^ T cells augmented macrophage *Il1b* mRNA expression **(Extended data Fig. 11e),** caspase-1 activity **(Fig. 5i, Extended data Fig. 11f),** and IL-1β release **(Fig. 5j).** The synergistic activity between macrophages and CD8^+^ T cells to produce IL-1β was not observed when isolated from naïve mice, suggesting that prior infection is necessary to prime the cells **(Extended data Fig. 11g)**. Moreover, IL-1β released into the supernatant was significantly reduced upon treatment with IFN-γ and TNF neutralizing Ab in the coculture system, confirming the role of IFN-γ and TNF in promoting IL-1β release by macrophages **(Fig. 5k)**. Consistently, we observed a similar decrease in BAL fluid IL-1β levels following treatment with either αCD8 **(Fig. 5l)**, or αIFN-γ + αTNF treatment **(Fig. 5m)**.

Since CD8^+^ T cells, monocyte-derived macrophages, and Krt8^hi^ progenitors accumulate within dysplastic areas after influenza infection, we tested whether macrophage-derived IL-1β is a negative regulator of AT2 to AT1 trans-differentiation. Using a 2D primary AT2 cell culture model known to induce spontaneous differentiation into AT1 cells through the transitional cell stage, we examined if conditioned media from cocultured CD8^+^ T cells and macrophages influenced AT2 trans-differentiation **(Fig. 5n)** (18, 53). AT1 cell marker expression was reduced upon exposure to conditioned media from CD8^+^ T cell-macrophage coculture compared to macrophages alone, which was rescued upon treatment with αIL-1β **(Fig. 5o)**. In contrast, transitional cell marker expression exhibited the opposite pattern, with increased levels within the coculture group and a dramatic reduction following αIL-1β treatment **(Fig. 5p)**. Conversely, rIL-1β treatment inhibited AT1 marker expression and promoted the expression transitional cell markers akin to previous reports (21). Collectively, our results suggest that exuberant CD8^+^ T cell-macrophage interactions promote chronic IL-1β release to inhibit AT2 cell trans-differentiation by arresting the cells in the transitional state.

### Therapeutic neutralization of IFN-γ and TNF, or IL-1β activity enhances alveolar regeneration and restores lung function

Our data thus far indicate a pathological role for CD8^+^ T cells persisting in human PASC-PF or post-viral lungs in an animal model, in the development of chronic pulmonary sequelae. Since depletion of CD8^+^ T cells is not a clinically feasible treatment strategy, we explored the therapeutic efficacy of neutralizing the cytokines that are effectors of the profibrotic CD8^+^ T cell. As expected, blocking IFN-γ and TNF activity in aged influenza-infected mice ameliorated fibrotic sequelae when compared to isotype controls **(Fig. 6a,b, Extended data Fig. 12a)**. Moreover, IFN-γ and TNF neutralization diminished Krt5^+^ and Krt8^hi^ areas of dysplastic repair **(Fig. 6c,d)** and enhanced alveolar regeneration as evidenced by the increased numbers of AT1 and AT2 cells **(Fig. 6c,e)**. The observed cellular changes were also reflected in physiological benefit, with improved lung function following treatment **(Fig. 6f, Extended data Fig. 12b,c)**.

**Fig. 6.**
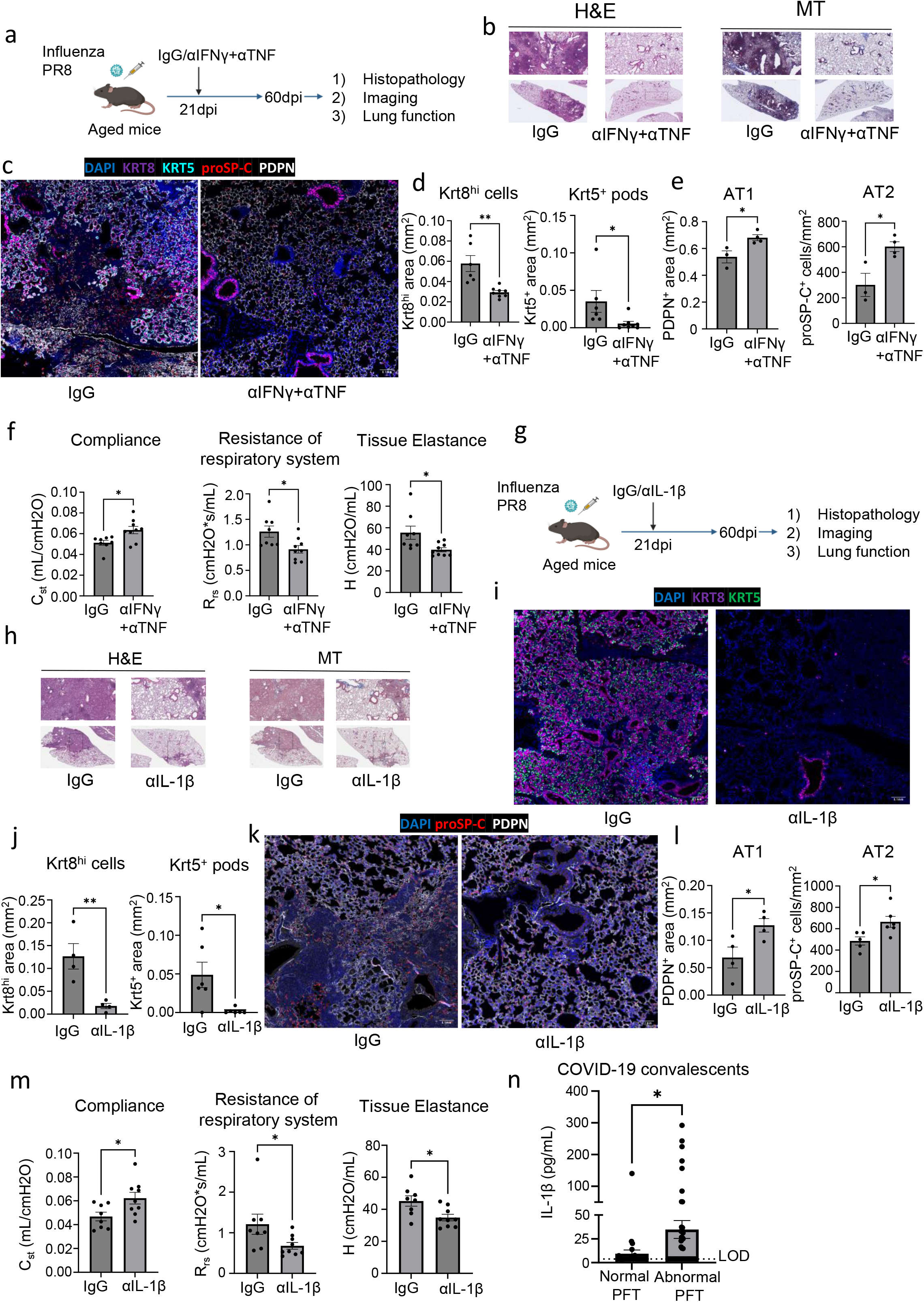
Therapeutic neutralization of IFN-γ and TNF, or IL-1β activity promotes alveolar regeneration and restores lung function. **a.** Experimental design for *in vivo* IFN-γ + TNF neutralization post influenza infection. **b.** Representative H&E and MT images of aged C57BL/6 lungs post influenza infection (42pi) treated with αIFN-γ + αTNF neutralizing Ab or control IgG Ab. **c.** Representative immunofluorescence images staining AT1 (PDPN^+^), AT2 (proSP-C^+^) and epithelial progenitors (Krt5^+^ and Krt8^hi^) in aged influenza-infected mice (42dpi) treated with αIFN-γ + αTNF neutralizing Ab or control IgG Ab. **d.** Quantification of Krt8^hi^ and Krt5^+^ area and **e.** AT1 (PDPN^+^) and AT2 (proSP-C^+^) cells in aged influenza-infected mice treated with αIFN-γ + αTNF neutralizing Ab or control IgG Ab (n= 3-6 IgG, 4-8 αIFN-γ + αTNF). **f.** Evaluation of static compliance (C_st_), resistance of the respiratory system (R_rs_), and tissue elastance (H) in aged influenza-infected mice treated with αIFN-γ + αTNF neutralizing Ab or control IgG Ab (n= 8 IgG, 10 αIFN-γ + αTNF). **g.** Experimental design for *in vivo* IL-1β blockade post influenza infection. **h.** Representative H&E and MT images of aged C57BL/6 lungs post influenza infection (42pi) treated with αIL-1β or control IgG Ab. **i.** Representative immunofluorescence images staining epithelial progenitors (Krt5^+^ and Krt8^hi^) and **k.** AT1 (PDPN^+^), AT2 (proSP-C^+^) in aged C57BL/6 mouse lungs post influenza infection (42dpi) treated with αIL-1β or control IgG Ab. **j.** Quantification of Krt8^hi^ and Krt5^+^ areas, and **l.** AT1 (PDPN^+^) and AT2 (proSP-C^+^) cells in aged influenza-infected lung sections treated with αIL-1β or control IgG Ab (n= 9 IgG, 10 αIL-1β). **m.** Evaluation of static compliance (C_st_), resistance of the respiratory system (R_rs_), and tissue elastance (H) in aged influenza-infected lung sections treated with αIL-1β neutralizing antibody or control IgG antibody (n= 9 IgG, 10 αIL-1β). **n.** Evaluation of IL-1β levels in plasma of COVID-19 convalescents with or without abnormal lung function. Data are expressed as mean ± SEM. Statistical analyses were conducted using a two-tailed unpaired t-test (d-f, j, l-n). *p < 0.05; **p < 0.01.

Next, we tested the efficacy of neutralizing IL-1β in influenza-infected aged mice and observed dramatic attenuation of lung fibrosis, which phenocopies the results of CD8^+^ T cell depletion, and neutralization of IFN-γ and TNF **(Fig. 6g,h, Extended data Fig. 12d)**. Improved alveolar regeneration was also observed, as evidenced by reduced Krt5^+^ and Krt8^hi^ areas and increased AT1 and AT2 cells **(Fig. 6i-l)**. Further confirming the therapeutic efficacy of IL-1β blockade post infection, we found improved lung function in Ab-treated mice **(Fig. 6m, Extended data Fig. 12e,f)**. Notably, improved outcomes following IL-1β neutralization were only seen in aged mice but not young mice, which is likely due to the sustained elevation of IL-1β observed exclusively in aged mice (**Extended data Fig. 12g,h).** Consistent with the observed improvement in outcomes and a recent study identifying increased BAL IL-1β levels in respiratory PASC (7), we found that circulating IL-1β levels were elevated in individuals exhibiting persistent abnormal pulmonary function compared to those that had fully recovered **(Fig. 6n),** suggesting that chronic IL-1β activity may impede the restoration of normal lung function after acute SARS-CoV-2 infection.

Given the efficacy of inhibiting IL-6 activity in treating acute COVID-19 (54, 55) as well as preclinical studies suggesting rescue of fibrotic disease after bleomycin injury (56–58), we also investigated the utility of IL-6 blockade in our model **(Extended data Fig. 13a)**. In contrast to the neutralization of IFN-γ and TNF or IL-1β, neutralizing IL-6 activity in the post-acute phase of infection did not improve the pathology or fibrotic sequelae post infection **(Extended data Fig. 13b,c)**. Similarly, no effect was observed on Krt8^hi^ and Krt5^+^ areas, alveolar regeneration, or pulmonary function **(Extended data Fig. 13d-g)**. Although Krt5^+^ progenitors were previously shown to migrate to the distal airspace in response to IL-6 signaling (15), our data indicates that their maintenance after the formation of pods in the alveolar epithelium is not dependent on IL-6 signaling.

Collectively, these data support the neutralization of IFN-γ and TNF, or IL-1β in the post-acute stage of viral infection as viable therapeutic options to augment alveolar regeneration and dampen fibrotic sequelae observed following respiratory viral infections (**Extended data Fig. 14**).

## DISCUSSION

Recent efforts have revealed several histopathological features conserved across various cohorts of respiratory PASC and PASC-PF patients including the prolonged reduction in alveolar epithelial cells (38), maintenance of Krt8^hi^ and Krt5^+^ dysplastic progenitors (5, 8), and the persistence of various immune cell populations in the lungs (9, 10, 59). Although immune-derived cues have recently been shown to influence lung repair (11, 21, 47, 50, 60), their exact interactions, if any, with the alveolar epithelium and role in post-viral fibrosis remain unexplored. Here, we used three independent cohorts of respiratory PASC to identify spatially defined microenvironments composed of a dysregulated immune-epithelial progenitor niche, a unique feature of PASC-PF but not acute COVID-19 or IPF lungs. With insights from patient samples, we established a clinically relevant animal model of post-viral fibrosis to demonstrate a causal role for exuberant proximal interactions between CD8^+^ T cells, macrophages, and epithelial progenitors in driving chronic tissue sequelae and fibrosis after viral pneumonia.

Understanding the mechanistic basis of respiratory PASC requires suitable animal models. Although SARS-CoV-2 MA-10 infection of C57BL/6, BALB/c, and K18-hACE2 mice initially caused severe alveolar pathology, minimal fibrosis was observed beyond acute disease. In contrast to previous studies suggesting aged BALB/c mice develop significant pulmonary inflammation and prolonged pathology after SARS-CoV-2 infection, our data indicate these murine SARS-CoV-2 infection models may not fully recapitulate the pathophysiology leading to PASC-PF in humans. A major deficiency of the SARS-CoV-2 mouse infection model is the absence of persistent Krt8^hi^ and Krt5^+^ areas of dysplastic repair – a hallmark of human PASC-PF. Moreover, CD8^+^ T cells, which are also enriched in human PASC-PF lungs to potentially promote the maintenance of Krt8^hi^ and Krt5^+^ areas, are not appreciably increased post SARS-CoV-2 infection in BALB/c or K18-hACE2 mice (9), which may be a reflection of a genetically programmed bias towards T_H_2 responses (61) and the lack of substantial acute lung injury, respectively. Thus, it is imperative to adopt clinically relevant animal models of SARS-CoV-2 post-viral fibrosis, validated by comparative analyses with human samples, to uncover the underlying mechanisms and identify therapeutic targets to treat PASC. In this study, we show that influenza infection in aged C57BL/6 mice induced chronic pulmonary sequelae that was maintained up to 250 days post infection and faithfully recapitulated the immunopathological features of human PASC-PF lungs.

Excessive infiltration and accumulation of profibrotic monocyte derived macrophages has been reported in the context of severe acute COVID-19, IPF, and PASC (12, 59, 62–64). Here, we elucidate a previously unknown role for pulmonary CD8^+^ T cells in impaired recovery and fibrotic remodeling in PASC-PF but not acute COVID-19 or IPF lungs. This distinction is likely the product of a dysregulated and protracted antiviral response, originally aimed towards the clearance of the virus and virus-infected cells (65). Typically, these pulmonary CD8^+^ T cells gradually contract with successful alveolar regeneration in individuals that successfully recover from acute COVID-19 (9). In human PASC lungs however, the long-term persistence of CD8^+^ T cells impairs recovery post infection and drives the development of fibrotic disease. It is unclear if the prolonged maintenance and activity of CD8^+^ T cells in lungs are a result of excessive TGF-β signaling reported to occur in PASC (8), chronic persistence of viral remnants (66–69), or other independent mechanisms. Nevertheless, the prominent spatiotemporal association between the epithelial progenitors and CD8^+^ T cells in the post-viral lung strongly suggests the production of T cell-recruitment and/or survival signals by the epithelium, whose identity remains unknown. Given the overlap in fibrogenic pathways, it has been proposed that PASC-PF may represent an intermediate state prior to potential progression towards IPF (8, 32). Whether CD8^+^ T cells primarily dictate the balance between functional recovery and PASC-PF post infection, or also play a pivotal role in the development of IPF is an open question (70).

By combining imaging and spatial transcriptomics modalities, we show that respiratory sequelae post viral infections is, at least in part, a result of chronic IL-1β signaling downstream of aberrant interactions between CD8^+^ T cells and monocyte-derived macrophages, mediated by IFN-γ and TNF. Although chronic IL-1β was found to impair AT2 trans-differentiation *in vitro*, the use of aged mice in our experiments posed a logistical challenge to directly test the effect of IL-1β on Krt8^hi^ and Krt5^+^ progenitors *in vivo* using transgenic mice. Thus, there exists a distinct possibility for IFN-γ and TNF as well as IL-1β and relevant downstream mediators to influence epithelial progenitor cell fate through their actions on other lung immune and non-immune cells to ultimately result in fibrotic remodeling. These data deepen our fundamental understanding of the role of the immune system in alveolar regeneration and post-viral lung disease. It is imperative to further extend this work and elucidate the role of other immunological players present within the post-viral lung and similarly understand the respective contributions of the mesenchyme and the endothelium in dictating the balance between functional and pathological lung repair. Additionally, in contrast to anti-viral (71) agents that require early intervention, we show that neutralization of IFN-γ and TNF, or IL-1β activity in the aftermath of infection can serve as “pro-repair strategies” to augment alveolar regeneration and dampen fibrotic sequelae. As PASC and other post-viral pathologies are typically diagnosed several weeks or months after the primary infection, it is critical to identify treatment approaches capable of effectively rescuing disease upon administration in the chronic phase. Our data strongly suggest that drugs such as Anakinra, an IL-1 receptor antagonist or Baricitinib, a JAK-inhibitor, which have already been granted emergency use authorization for acute COVID-19 (72, 73), may serve as promising candidates to address this unmet need and treat ongoing respiratory PASC in the clinic.

## Acknowledgements

We thank the Lung Institute BioBank study staff at Cedars-Sinai Medical Center for providing human lung specimens and patients’ clinical data. We thank Drs. Andrew Vaughan and Rachel Zemans for protocols to isolate and culture AT2 cells. Schematics in the manuscript were created with BioRender.com. Data for this manuscript were generated using the Flow Cytometry and Research Histology Core Facilities at the University of Virginia. 10X Visium spatial transcriptomics on the human and mouse tissues were performed in the Applied Genomics, Computation and Translational Core at Cedars-Sinai Medical Center and the Advanced Genomics Core at the University of Michigan, respectively.

The study was in part supported by the US National Institutes of Health grants AI147394, AG069264, AI112844, HL170961, AI176171 and AI154598 to J.S, R01HL132287, R01HL167202, and R01HL132177 to Y.M.S, and F31HL170746 and T32AI007496 to H.N.

## Author contribution

H.N. & J.S. conceived the overall project. H.N. I.S.C., W.Q., S.H., T. P., C.Z., P.C., and J.S. designed the experimental strategy and analyzed data. C.L., N.G., Y.W., X.W., Y.M.S., E.F, G.S., J.T., C.Y., L.M., G.C., A.K., A.C., S.P.Y., R.H., G.B., M.A., A.S.C., P.P., Y.M.S., and J.W., performed experiments, analyzed data, or contributed critical reagents to the study. H.N. & J.S. wrote the original draft. All authors read, edited, and approved the final manuscript.

## METHODS

### Ethics and biosafety

All aspects of this study were approved by the Institutional Review Board Committee at Cedars-Sinai Medical Center (IRB# Pro00035409) and the University of Virginia (IRB# 13166). Work related to SARS-CoV-2 was performed in animal biosafety level 3 (ABSL-3) facilities at the University of Virginia and influenza-related experiments were performed in animal biosafety level 2 (ABSL-2) facilities at the Mayo Clinic and the University of Virginia.

### Cells, viruses, and mice

African green monkey kidney cell line Vero E6 (ATCC CRL-1587) were maintained in Dulbecco modified Eagle medium (DMEM) supplemented with 10% fetal bovine serum (FBS), along with 1% of penicillin-streptomycin (P/S) and L-glutamine at 37°C in 5% CO_2_. The SARS-CoV-2 mouse-adapted strain (MA-10) was kindly provided by Dr. Barbara J Mann (University of Virginia School of Medicine). The virus was passaged in Vero E6 cells, and the titer was determined by plaque assay using Vero E6 cells.

Young (8–10-week-old) and aged (84-week-old) female C57BL/6J mice, and 15-week-old female BALB/cJ mice were purchased from The Jackson Laboratory (JAX). K18-hACE2 (catalog no. 034860) transgenic mice as well as WT C57BL/6 (catalog no. 000664) mice were purchased from the Jackson Laboratory and bred in-house. Aged mice were received at 20 to 21 months of age from the National Institutes of Aging and all mice were maintained in the facility for at least 1 month before infection. All mice were housed in a specific pathogen–free environment and used under conditions fully reviewed and approved by the institutional animal care and use committee guidelines at the Mayo Clinic (Rochester, MN) and University of Virginia (Charlottesville, VA).

For primary influenza virus infection, influenza A/PR8/34 strain [75 plaque-forming units (PFU) per mouse] was diluted in fetal bovine serum (FBS)–free Dulbecco’s modified Eagle’s medium (DMEM) (Corning) on ice and inoculated in anesthetized mice through intranasal route as described previously (42). For SARS-CoV-2 infections, mice were infected with 5 x 10^4^ PFU for C57/BL6 and 1000PFU for BALB/c of mouse-adapted (MA-10) virus or 200PFU of USA-WA1/2020 for K18-hACE2, intranasally under anesthesia as described previously. Infected mice were monitored daily for weight loss and clinical signs of disease for 2 weeks, following once a week for the duration of the experiments. The mortality rate of mice calculated as “dead” were either found dead in cage or were euthanized as mice reached 70% of their starting body weight which is the defined humane endpoint in accordance with the respective institutional animal protocols. At the designated endpoint, mice were humanely euthanized by ketamine/xylazine overdose and subsequent cervical dislocation.

### *In vivo* drug treatment

For depletion of CD8^+^ T cells and neutralization of cytokines, starting at 21dpi, mice were intraperitoneally administered 20μg or 500μg of anti-CD8 Ab (53-6.7 and 2.43; BioXCell), anti-IL-1β (B122; BioXCell), anti-IFNγ (XMG1.2; BioXCell), anti-TNFα (XT3.11; BioXCell) or isotype control immunoglobulin G (IgG) in 200μL of phosphate-buffered saline (PBS) once a week. Mice were euthanized 3 days after the last treatment.

### Evaluation of respiratory mechanics and lung function

Lung function measurements using forced oscillation technique and the resulting parameters have been previously described (74). In brief, animals were anesthetized with an overdose of ketamine/xylazine (100 and 10mg/kg intraperitoneally) and tracheostomized with a blunt 18-gauge canula (typical resistance of 0.18 cmH_2_O s/mL), which was secured in place with a nylon suture. Animals were connected to the computer-controlled piston (SCIREQ flexiVent), and forced oscillation mechanics were performed under tidal breathing conditions described in (74) with a positive-end expiratory pressure of 3 cm H_2_O. The measurements were repeated following thorough recruitment of closed airways (two maneuvers rapidly delivering total lung capacity of air and sustaining the required pressure for several seconds, mimicking holding of a deep breath). Each animal’s basal conditions were normalized to their own maximal capacity. Measurement of these parameters before and after lung inflation allows for determination of large and small airway dysfunction under tidal (baseline) breathing conditions. Only measurements that satisfied the constant-phase model fits were used (>90% threshold determined by software). After this procedure, mice had a heart rate of ∼60 beats per minute, indicating that measurements were done on live individuals.

### Human lung tissue specimens

Human lung samples were obtained from patients enrolled in the IRB-approved Lung Institute BioBank (LIBB) study at Cedars-Sinai Medical Center, Los Angeles, CA and at the University of Virginia, Charlottesville, VA. All participants or their legal representatives provided informed written consent. Lung tissues were processed within 24 hours after surgical removal. Specifically, the lung tissues were cut and immediately fixed in 10% normal-buffered formalin for 24 hours before tissue processing using the HistoCore PEARL – Tissue Processor, Leica Biosystem, Deer Park, IL, and embedded in paraffin for histological studies. The formalin-fixed paraffin-embedded cassettes were appropriately stored at room temperature until further sectioning.

### Mouse tissue processing and flow cytometric analysis

Animals were injected intravenously with 2μg of CD45 or CD90.2 Ab labeled with various fluorochromes. Two minutes after injection, animals were euthanized with an overdose of ketamine/xylazine and processed 3 min later. After euthanasia, the right ventricle of the heart was gently perfused with chilled 1X PBS (10 mL). The right lobes of the lungs were collected in 5 mL of digestion buffer (90% DMEM and 10% PBS and calcium with type 2 collagenase (180 U/ml) (Worthington) and DNase (15μg/ml) (Sigma-Aldrich) additives). Tissues were digested at 37°C for 1 hour followed by disruption using gentleMACS tissue dissociator (Miltenyi). Single-cell suspension was obtained by hypotonic lysis of red blood cells in ammonium-chloride-potassium buffer and filtration through a 70μm mesh. Cells were washed with FACS buffer (2% of FBS and 0.1% of NaN_3_ in PBS) and FC-γ receptors were blocked with anti-CD16/32 (2.4G2). Surface staining was performed by antibody (details provided in **Supplementary Table 2**) incubation for 30 min at 4°C in the dark. After PBS wash, cells were resuspended with Zombie-dye (Biolegend) and incubated at RT for 15 min. For IL-1β staining, cells were incubated with monesine (Biolegend) for 5 hours at 37°C and then stained with surface markers antibodies. After washing with FACS buffer, cells were fixed with fixation buffer (Biolegend) and permeabilized with intracellular staining permeabilization wash buffer (Biolegend). The cells were then stained with anti-IL-1β at RT for 1hr and samples were acquired on an Attune NxT (Life Technologies). The data were analyzed with FlowJo software (Tree Star).

### Caspase 1 FLICA analysis

Caspase 1 was detected by FAM-FLICA Caspase-1 Assay Kit (ImmunoChemistry Technologies) according to the manufacturer’s instructions. Briefly, lung single cells or macrophages from *in vitro* culture were stained with fluorochrome-conjugated Ab cocktail for cell surface markers. After staining, cells were incubated with FLICA for 30min at 37°C, washed and detected by flow cytometry.

### Cell isolation and *ex vivo* co-culture

To isolate myeloid and CD8^+^ T cells, single cell suspension of the lung was generated from influenza infected mice as described above and labeled and enriched with CD11c and CD11b or CD8 microbeads (Miltenyi Biotec, Auburn, CA) according to the manufacturer’s instructions. Purified myeloid cells were seeded in 96-well (200,000 per well) plates and incubated for 2 hours at 37°C 5% CO_2_ to facilitate attachment. Wells were washed with 1X PBS to select for macrophages, following which selected wells were seeded with naïve or memory CD8^+^ T cells (40,000 per well) and further incubated for 16-18 hours. Supernatant and cell pellets were collected for cytokine measurement and gene expression analyses, respectively.

### AT2 cell isolation and culture

AT2 cells were isolated from naïve mice as previously described (28, 53, 75). Briefly, mouse lungs were perfused with chilled PBS and intratracheally instilled with 1mL of dispase II (15U/mL, Roche), tying off the trachea and cutting away the lobes from the mainstem bronchi. Lungs were incubated in 4mL of 15U/mL dispase II for 45min while shaking at room temperature, followed by mechanical dissociation with an 18G needle. Following passage through a 100μm filter, lungs underwent 10min of DNase I digestion (50μg/mL) and filtered through 70μm filter prior to RBC lysis. Single-cell suspensions were subject to CD45 depletion using microbeads (Miltenyi), incubated with anti-FcgRIII/II (Fc block) and stained with CD45, EpCAM, MHC-II, and viability die (see antibody details in **Supplementary Table 2**). Fluorescence assisted cell sorting was performed on the BD Influx cell sorter to isolate AT2 cells as described previously (75) and collected in 500μL DMEM + 40% FBS + 2% P/S. Sorted AT2 cells (200,000 per well) were plated in a 96-well plate in DMEM/F12 + 10% FBS and cultured at 37°C, 5% CO_2_ for 3 days prior to harvest. For *in vitro* neutralization anti-IL-1β (B122; BioXCell) was used at 10μg/mL and for stimulation, IL-1β (Peprotech) was used at 20ng/mL.

### RNA isolation and real time-quantitative polymerase chain reaction (RT-qPCR)

Cells were lysed in Buffer RLT and RNA was purified using the RNeasy Plus Mini Kit (Qiagen) according to the manufacturer’s instructions. Random primers (Invitrogen) and MMLV reverse transcriptase (Invitrogen) were used to synthesize first-strand complementary DNAs (cDNAs) from equivalent amounts of RNA from each sample. cDNA was used for real-time PCR with Fast SYBR Green PCR Master Mix (Applied Biosystems). Real-time PCR was conducted on QuantStudio 3 (Applied Bioscience). Data were generated with the comparative threshold cycle (Δ*C*t) method by normalizing to hypoxanthine-guanine phosphoribosyltransferase (HPRT) transcripts in each sample as reported previously (76).

### COVID-19 convalescents cohort

Peripheral blood specimens were obtained from patients presenting to the University of Virginia Post-COVID-19 clinic. All participants provided written informed consent. Pulmonary function testing (PFT: spirometry, lung volume testing, diffusing capacity of the lungs for carbon monoxide [DLCO]) at the time of blood draws was used to define normal and abnormal lung function.

### IL-1β cytokine evaluation

Plasma from COVID-19 convalescents, supernatant from *ex vivo* culture and murine BAL fluid following flushing of airway 3X with 600uL of sterile PBS were used to quantify IL-1β levels by ELISA as per manufacturer’s instructions (R&D systems). Samples were first concentrated 5X (BAL) and 2X (cell culture supernatant) using Microcon-10kDa centrifugal filters (Millipore Sigma).

### Histopathological analysis for fibrosis

Mouse lungs were routinely perfused with ice cold 1X PBS and the left lung was inflated and fixed in 10% formalin, embedded in paraffin, and cut into 5μm thick sections. Sections were stained with hematoxylin eosin and Masson’s trichrome. All sections were studied using light microscopy. The severity of fibrosis was semi-quantitatively assessed by blinded reviewers according to a modified Ashcroft scale (77) and scored between 0 to 8 by examining randomly chosen fields.

### Immunofluorescence

Lung tissue sections (5μm) were deparaffinized in xylene and rehydrated. Heat-induced antigen retrieval was performed using 1X Agilent Dako target retrieval solution (pH 9) in a steamer for 20min (mouse lungs) or 45min (human lungs), followed by blocking and surface staining. For intracellular targets, tissues were permeabilized with 0.5% Triton-X 0.05% Tween20 for 1 hour at room temperature. Sections were stained with primary antibodies as listed in **Supplementary Table 2** overnight at 4°C. Subsequently, samples were washed and incubated with fluorescent secondary antibodies as listed in **Supplementary Table 2** for 2 hours at room temperature. Sections were counterstained with DAPI (1:1000, ThermoFisher Scientific) for 3 minutes and mounted using ProLong Diamond Antifade mountant (ThermoFisher Scientific). After 24 hours of curing at room temperature, images were acquired using a 10X or 20X objective on the Olympus BX63 fluorescent microscope and pseudocolours were assigned for visualization. For each lung section, images were taken in at least 10-12 random areas in the distal lung for quantification. All images were further processed using ImageJ Fiji, OlyVIA, and/or QuPath software (78). Quantitative analysis of stitched immunofluorescence images was performed using QuPath v0.4.1, where tissue-specific expression of Krt8^hi^ was used to annotate areas of dysplastic repair and discriminate them from the remaining intact human or mouse lung tissue. CD8^+^ T cells and macrophages were detected using CD8α or CX3CR1 (mouse)/CD68 (human) signals, respectively. The abundance and distribution of the respective cells of interest were quantified to plot distance from the annotation (Krt8^hi^). Density maps were further generated to identify hotspots of the cells across the tissue with respect to the annotations. The same parameters for annotation and cell-type classification were applied to all samples from the same experiment.

### Spatial Transcriptomics analyses

Spatial transcriptomics (ST) data (generated from 10x Visium platform) was pre-processed using the spaceranger package (v2.0, genome version mm10). The R package Seurat (v4.3.0) was used for quality control (QC) and basic analyses. Only high-quality spots with sufficient gene coverage (>=2000) were retained for downstream analysis. Like scRNA-seq data, the spatial expression data was first normalized to a log scale using SCTransform method (v0.3.5). The top 2000 Highly variable genes were then identified based on the variance of expression across spots for the principal component analysis (PCA) input. UMAP embeddings were generated for visualization at the reduced dimensionality (top 30 PCs). Spots were clustered with a shared nearest neighbor (SNN) modularity optimization-based clustering algorithm (built in Seurat). Samples were integrated and batch-effects were removed using the Harmony package (v0.1). In addition to the spatial spot view of the expression pattern, we used UMAP to visualize the expression pattern of spots associated with the identified cell types in a lower dimension using the visualization pipeline in the Seurat package. Marker genes (DEG) detection among different groups of spots was also performed with Seurat (FindMarker function), and the marker gene lists were further submitted to the GSEA pre-rank test for the functional annotation analysis.

We determined each spot’s potential cell type composition based on the following protocol: 1. We defined AE spots based on the k-means clustering analysis (using knowledge based key genes from different cell types). 2. For non-AE spots, we used the average expression of Krt5, Krt8, Krt17, Cldn4, and Trp53 to estimate a Krt score, and similarly, average expression of Cd8a, Cd8b1, Itgae, Trbc1, Trbc2, Sell, Ccl5, Cd69, and Cd3d for Cd8 score. We set −0.4 as cutoff for both the Cd8 and Krt score: spots with Krt/Cd8 score >-0.4 are assigned as Krt/Cd8 associated spot, and assigned blue/red color, respectively in the spatial map. In this way, a spot can be Krt spot and Cd8 spot at the same time, and we use purple color to represent this category. For a given gene list, captured the expression of relevant genes in Krt spots and AE spots and compared the distribution of the expression index. The significance of the difference was estimated by Wilcoxon rank sum test. The p-values were then log_10_ transformed and displayed as bar plot.

For mouse ST data, we determined each spot’s potential cell type composition based on the following protocol: 1. We define alveolar epithelium (AE) spots based on the k-means clustering analysis (using knowledge-based key genes from different cell types). 2. For non-AE spots, we use the average expression of Krt marker gene list **(Supplementary Table 4)** to estimate a Krt score, and similarly, average expression of Cd8 marker gene list **(Supplementary Table 4)** for Cd8 score. We set −0.4 as cutoff for both the Cd8 and Krt score: spots with Krt/Cd8 score > −0.4 are assigned as Krt/Cd8 associated spot, and assigned blue/red color, respectively in the spatial map. Thus, a spot can exhibit Krt and Cd8 signature at the same time, and we use purple color to represent this category. For a given gene list, we captured the expression of relevant genes to compare the distribution of their expression index between Krt spots and AE spots. The significance of the difference was estimated by Wilcoxon rank sum test. The p-values were then log10 transformed and displayed as barplot. Similar to the Cd8 score and Krt score, we also defined some other cell type scores based on our knowledge-based marker gene list **(Supplementary Table 4)**. We further utilized these scores to estimate the association between cell types in a specific slide or global level (by Pearson correlation coefficient). A pair of cell types with a higher Pearson correlation coefficient between their cell type score indicates a higher likelihood of their co-occurrence within the same spots.

For human ST data, most of the analysis was done with similar methods as mouse data. Additionally, we split spots by the expression of KRT8 and AGER using the cutoff of normalized expression index greater than 4. The spots with “KRT8 > 4 and AGER <= 4” were defined as “KRT8-only spots”, while the spots with “KRT8 <= 4 and AGER > 4” were defined as “AGER-only spots.” These two spot sets were collected to compare the difference in cell type scores, key gene expression, and pathway enrichment between them. In detail, we compared the percentage of ON spots (given a cell type score or gene expression index passing the cutoff) between the two groups (e.g., KRT17-only spots vs. AGER-only spots, or covid spots vs. control spots), and displayed the results with bar plots. A similar analysis was applied to examine the enrichment of relevant signaling pathways. Note that the cutoffs of normalized single gene expression were set to 0.8. The cell type score (e.g., CD8, KRT, AE, MDM, and AM scores) was set to 1.2 for human data as the distribution of marker gene expression varied between different species and biological samples (i.e., human lung covid samples). Detailed marker gene lists for cell types and pathways can be found in **(Supplementary Table 4)**.

### Statistical analyses

Quantitative data are presented as means ± SEM. Unpaired two-tailed Student’s *t* test (two-tailed, unequal variance) was used to determine statistical significance with Prism software (GraphPad) for two-group comparison. For multiple groups, analysis of variance (ANOVA) corrected for multiple comparisons was used when appropriate (GraphPad). Log-rank (Mantel-Cox) test was used for survival curve comparison and multiple *t* test was used to analyze differences in weight loss. We considered *P* < 0.05 as significant in all statistical tests and denoted within figures as a * for each order of magnitude *P* value.

---

**Extended data Fig.1.**
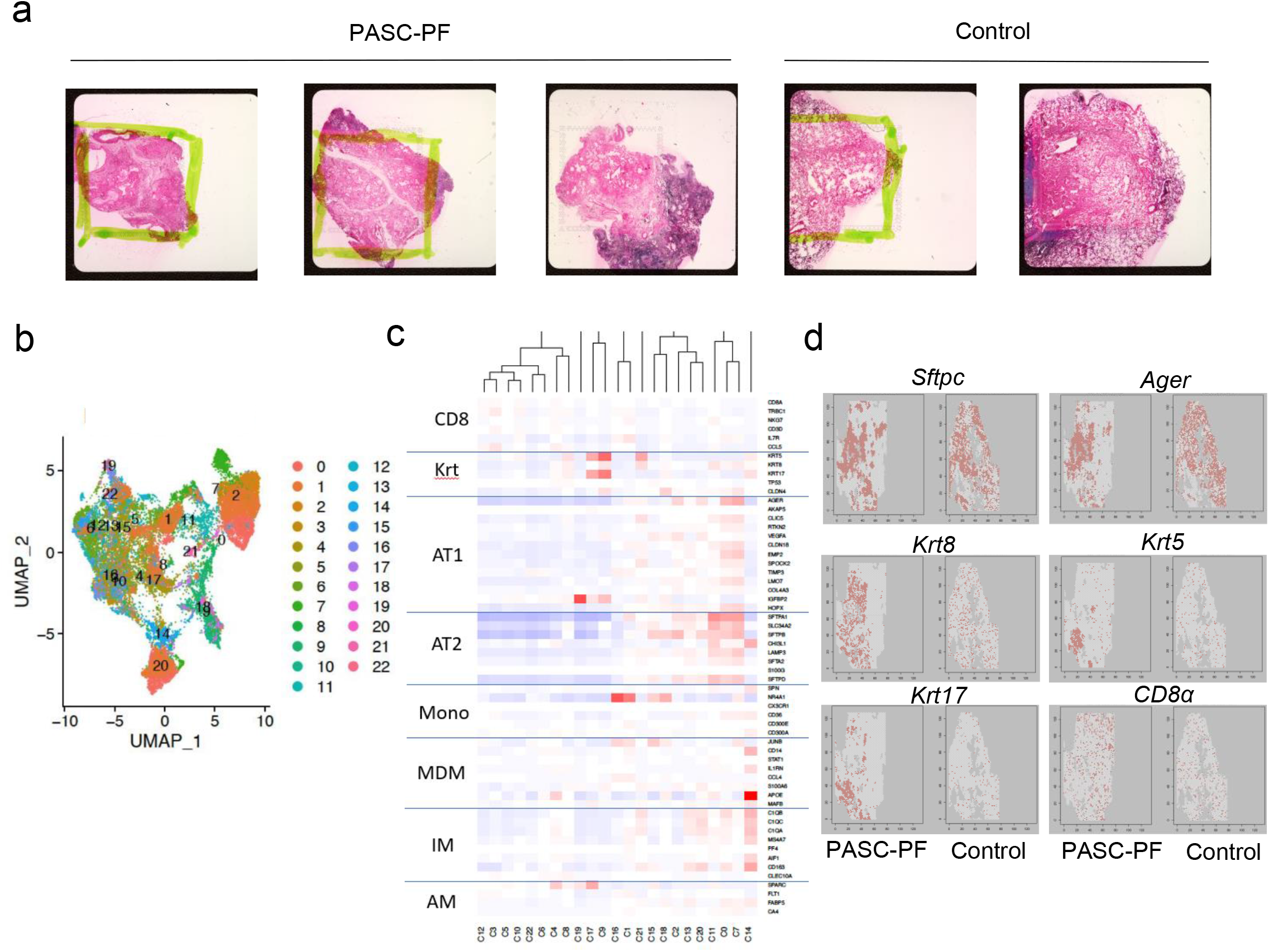
Spatial transcriptomics of human PASC-PF lungs reveal persistent defects in alveolar regeneration and chronic inflammation. **(a)** Representative H&E images of human control (n=2) and PASC-PF (n=3) lungs sections that were mounted onto the 10X Visium slide. **(b)** UMAP visualization of spatial transcriptomics data from human control and PASC-PF lungs. **(c)** Heatmap of key marker gene expression across all identified clusters of spots. **(d)** Spatial gene expression maps of epithelial and immune cell markers in control and PASC-PF lungs.

**Extended data Fig.2.**
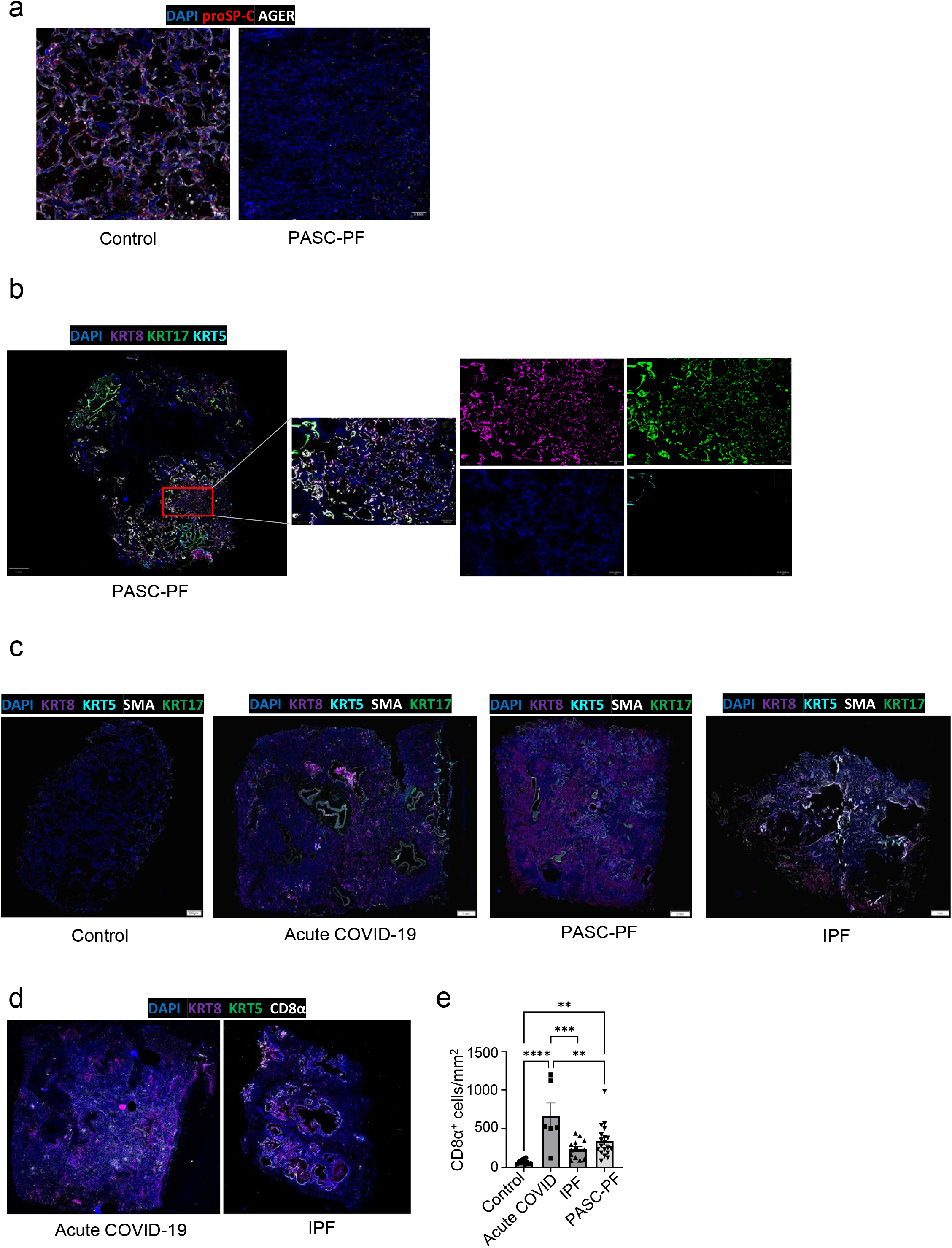
PASC-PF lungs exhibit hallmarks of persistent dysplastic repair and inflammation. **(a)** Representative immunofluorescence images staining alveolar epithelial cell markers (AT1 – AGER; AT2 – proSP-C) in control and PASC-PF lung sections. **(b)** Representative immunofluorescence image of PASC-PF lungs staining for dysplastic epithelial progenitors (Krt5, Krt8, Krt17), with higher magnification inset showing independent channels. **(c)** Representative immunofluorescence images staining for myofibroblasts (αSMA) and aberrant epithelial progenitors (Krt5, Krt17, Krt8) in human control, acute COVID-19, PASC-PF and IPF lung sections. **(d)** Representative immunofluorescence images staining CD8^+^ T cells and epithelial progenitors in acute COVID-19 and idiopathic pulmonary fibrosis (IPF) lung sections. **(e)** Quantification of CD8^+^ T cell number in control, acute COVID-19, PASC-PF, and IPF lung sections (n = 11 control, 6 Acute COVID-19, 19 PASC-PF, 13 IPF). Data are expressed as mean ± SEM. Statistical analyses were conducted using an ordinary one-way ANOVA (o). *p < 0.05; **p < 0.01; ***p < 0.001; ****p < 0.0001.

**Extended data Fig.3.**
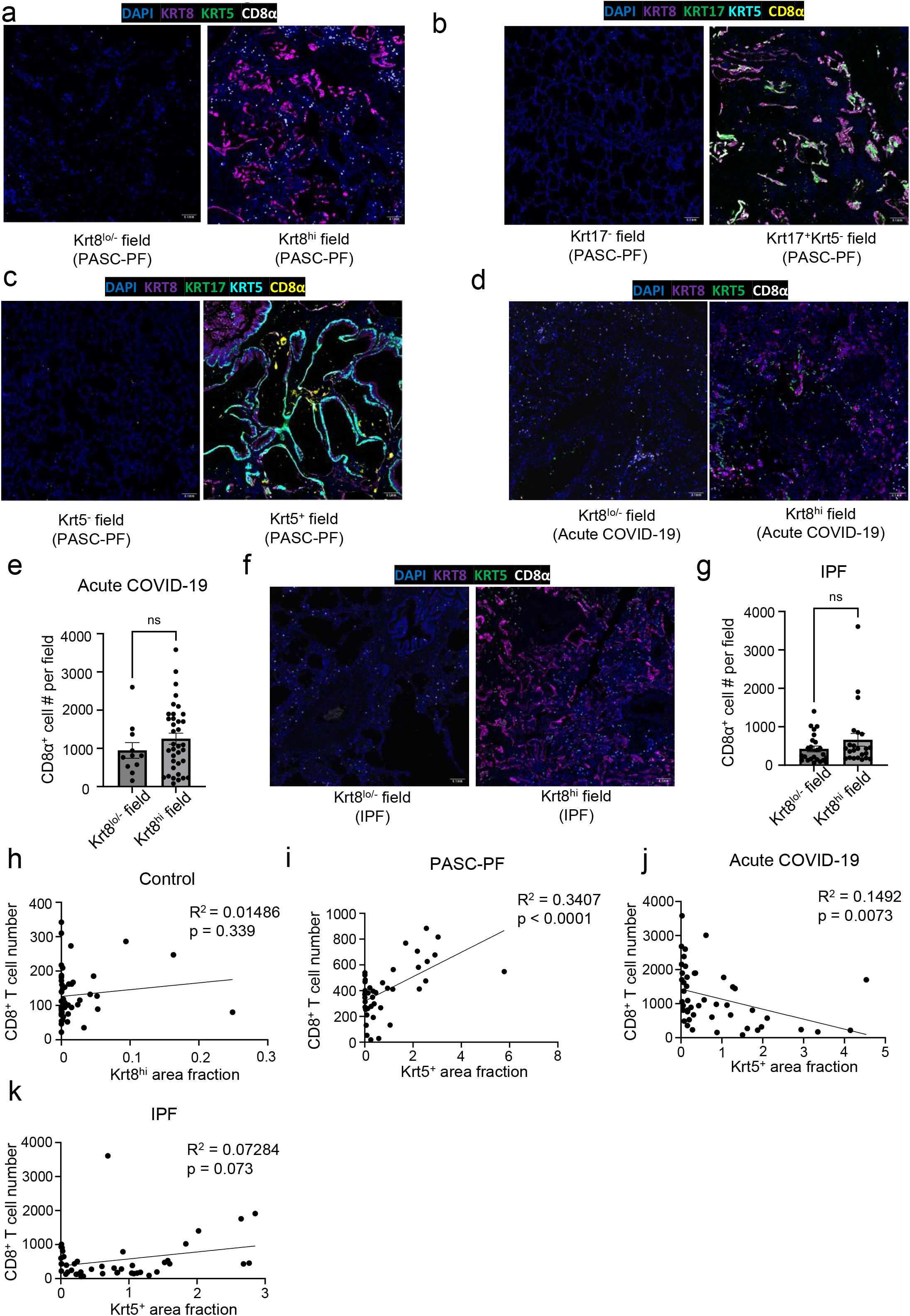
Characterization of immune-epithelial progenitor interactions in human PASC-PF, acute COVID-19 and IPF lungs. (**a)** Representative immunofluorescence images staining CD8^+^ T cells in Krt8^-/lo^ and Krt8^hi^ areas within PASC-PF lungs. **(b)** Representative immunofluorescence images staining CD8^+^ T cells in Krt17^-^ and Krt17^+^Krt5^-^ areas within PASC-PF lungs. **(c)** Representative immunofluorescence images staining CD8^+^ T cells in Krt5^-^ and Krt5^+^ areas within PASC-PF lungs. **(d)** Representative immunofluorescence images staining CD8^+^ T cells in Krt8^-/lo^ and Krt8^hi^ areas within acute COVID-19 lungs. **(e)** Quantification of CD8^+^ T cells in Krt8^-/lo^ and Krt8^hi^ areas within acute COVID-19 lungs. (n=8) **(f)** Representative immunofluorescence images staining CD8^+^ T cells in Krt8^-/lo^ and Krt8^hi^ areas within IPF lungs. **(g)** Quantification of CD8^+^ T cells in Krt8^-/lo^ and Krt8^hi^ areas within IPF lungs. (n=8) **(h)** Simple linear regression of CD8^+^ T cell number and Krt5^+^ area fraction in control lungs. **(i)** Simple linear regression of CD8^+^ T cell number and Krt5^+^ area fraction in PASC-PF, **(j)** acute COVID-19, and **(k)** IPF lung sections. Data are expressed as mean ± SEM. Statistical analyses were conducted using a two-tailed unpaired t-test. ****p<0.0001.

**Extended data Fig.4.**
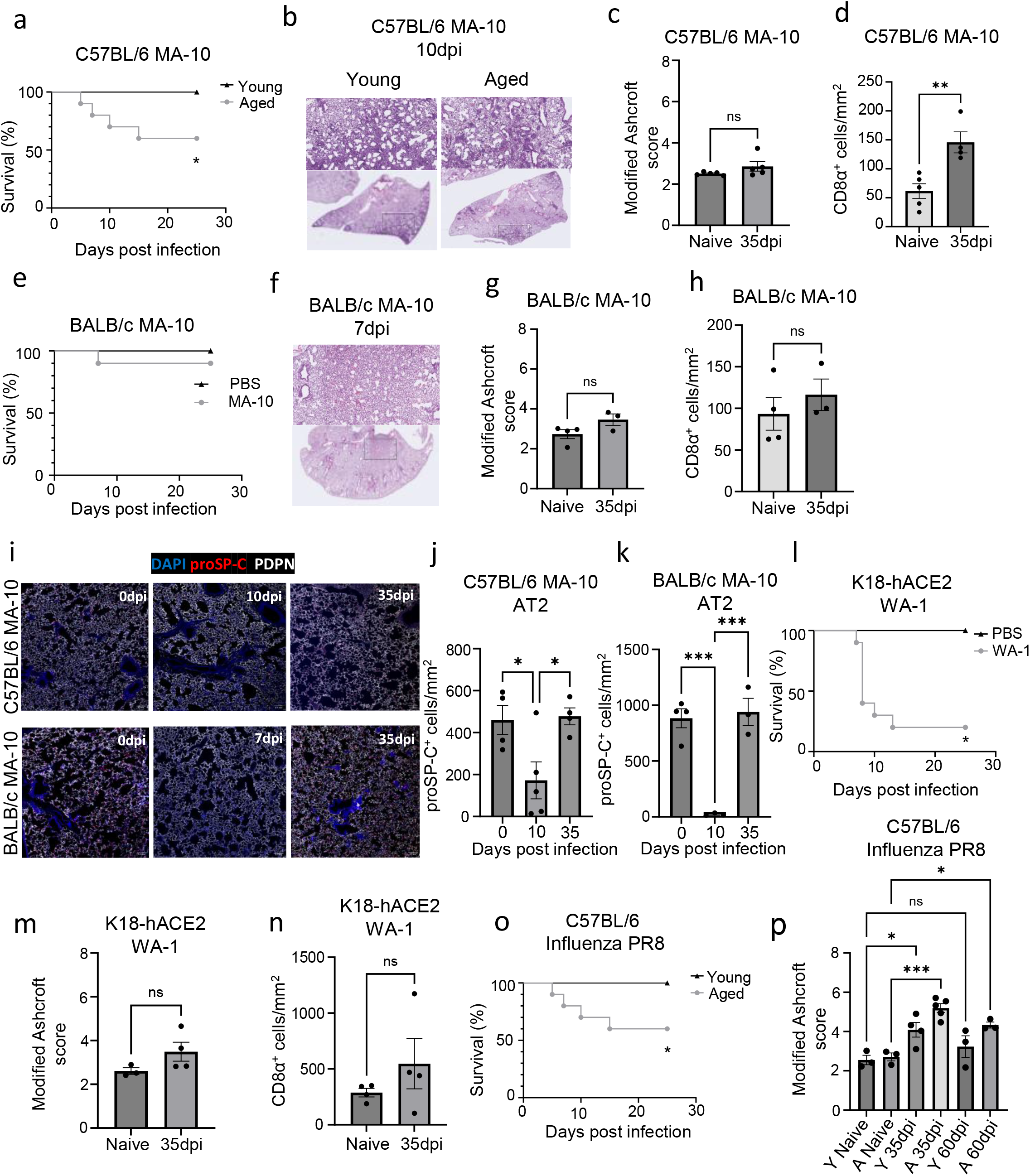
Aged mice exhibit increased severity during acute SARS-CoV-2 and influenza virus infection. **(a)** Survival data of young and aged C57BL/6 following SARS-CoV-2 MA-10 infection. (**b)** Representative H&E images of young and aged C57BL/6 mice from the acute (10dpi) phase of SARS-CoV-2 MA-10 infection. (**c)** Evaluation of fibrotic disease in naïve and SARS-CoV-2 MA-10 infected (35dpi) lungs of aged C57BL/6 mice by modified Ashcroft score. (n = 5 naïve, 5 MA-10) **(d)** Quantification of CD8^+^ T cell number from immunofluorescence images in naïve and SARS-CoV-2 MA-10 infected (35dpi) aged C57BL/6 mice. (n = 5 naïve, 4 MA-10) (**e)** Survival data of aged BALB/c following SARS-CoV-2 MA-10 infection. **(f)** Representative H&E images of aged BALB/c mice from the acute (7dpi) phase of SARS-CoV-2 MA-10 infection. (**g)** Evaluation of fibrotic disease in naïve and SARS-CoV-2 MA-10 infected (35dpi) lungs of aged BALB/c mice by modified Ashcroft score. (n = 4 naïve, 3 MA-10) **(h)** Quantification of CD8^+^ T cell number from immunofluorescence images in naïve and SARS-CoV-2 MA-10 infected (35dpi) aged BALB/c mice. (n = 4 naïve, 3 MA-10) **(i)** Representative immunofluorescence images of aged C57BL/6 and BALB/c mouse lungs infected with SARS-CoV-2 MA-10, staining alveolar epithelial cell markers (AT1-PDPN; AT2 – proSP-C). **(j)** Quantification of proSP-C^+^ AT2 cells in aged C57BL/6 and **(k)** BALB/c mouse lungs post SARS-CoV-2 MA-10 infection. (n= 3-5 per time point) **(l)** Survival data of aged K18-hACE2 mice following SARS-CoV-2 WA-1 virus infection. **(m)** Evaluation of fibrotic disease in naïve and SARS-CoV-2 WA-1 infected (35dpi) lungs of aged K18-hACE2 mice by modified Ashcroft score. (n = 3 naïve, 4 WA-1) **(n)** Quantification of CD8^+^ T cell number from immunofluorescence images in naïve and SARS-CoV-2 WA-1 infected (35dpi) aged K18-hACE2 mice. (n = 4 naïve, 4 WA-1) **(o)** Survival data of young and aged C57BL/6 following PR8 influenza virus infection. **(p)** Evaluation of fibrotic disease in naïve and PR8 influenza-infected (35dpi & 60dpi) lungs of young and aged C57BL/6 mice by modified Ashcroft score (n=3 per timepoint). Data are expressed as mean ± SEM. Statistical analyses were conducted using a two-tailed unpaired t-test (c,d,g,h,m,n), log-rank test (a,e,l,o), and an ordinary one-way ANOVA (j,k,p). *p < 0.05; **p < 0.01; ***p < 0.001.

**Extended data Fig.5.**
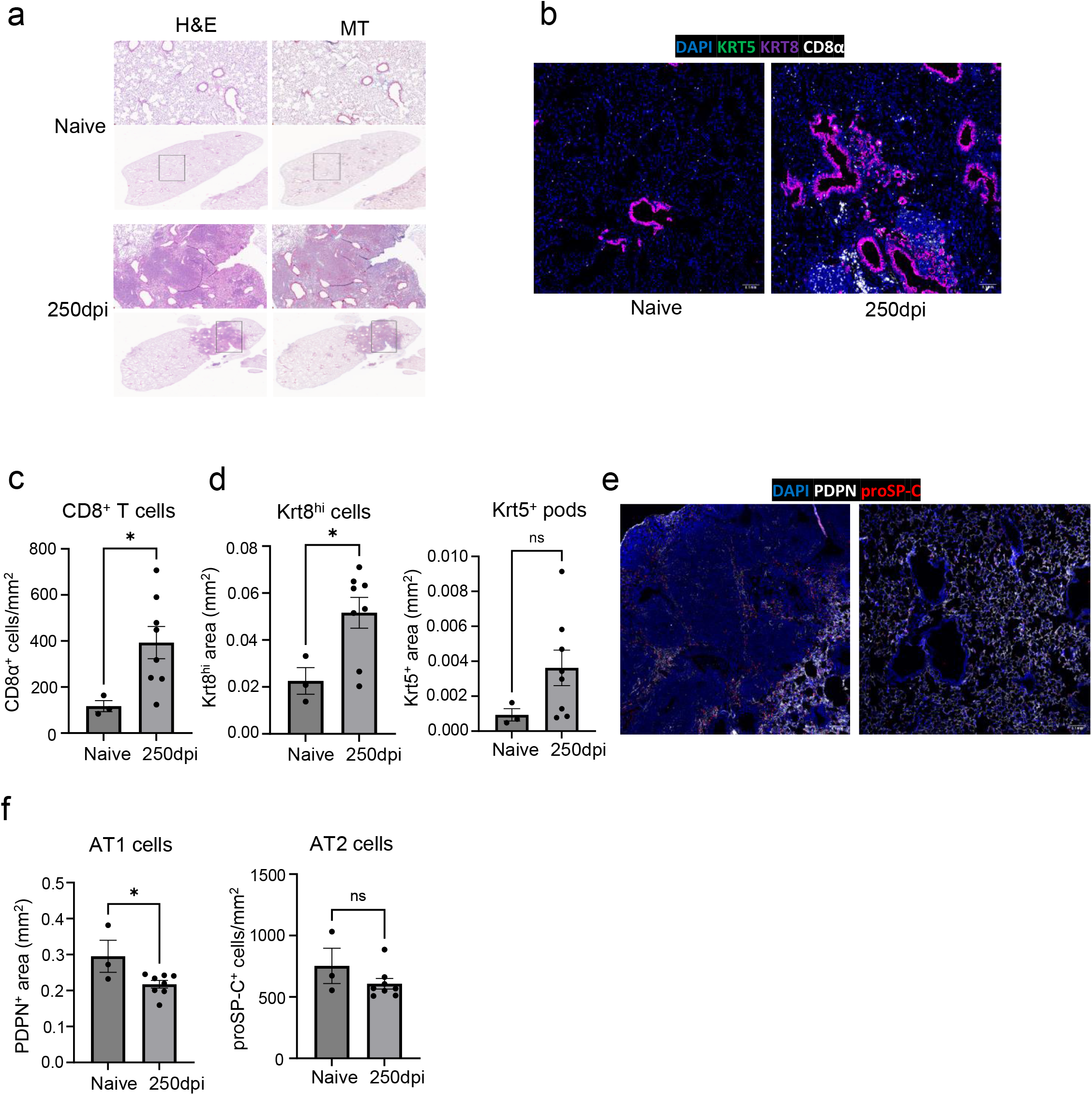
Aged influenza-infected mice exhibit chronic pulmonary pathology up to 8 months post infection. **(a)** Representative H&E and MT images of naive and influenza-infected (250dpi) lungs of aged C57BL/6 mouse lungs. **(b)** Representative immunofluorescence images of naive and influenza-infected (250dpi) aged C57BL/6 mouse lungs staining for dysplastic epithelial progenitors (Krt5 & Krt8) and CD8 T cells (CD8α). **(c)** Quantification of CD8^+^ T cells, **(d)** Krt8^hi^ and Krt5^+^ areas in naïve influenza-infected (250dpi) aged C57BL/6 mouse lungs. (n = 3 naïve, 8 infected) **(e)** Representative immunofluorescence images of naive and influenza-infected (250dpi) aged C57BL/6 mouse lungs staining for alveolar epithelial cell (AT1-PDPN; AT2 – proSP-C) markers. **(f)** Quantification of PDPN^+^ AT1 cells and proSP-C^+^ AT2 cells in naïve influenza-infected (250dpi) aged C57BL/6 mouse lungs (n = 3 naïve, 8 infected). Data are expressed as mean ± SEM. Statistical analyses were conducted using a two-tailed unpaired t-test. *p < 0.05.

**Extended data Fig.6.**
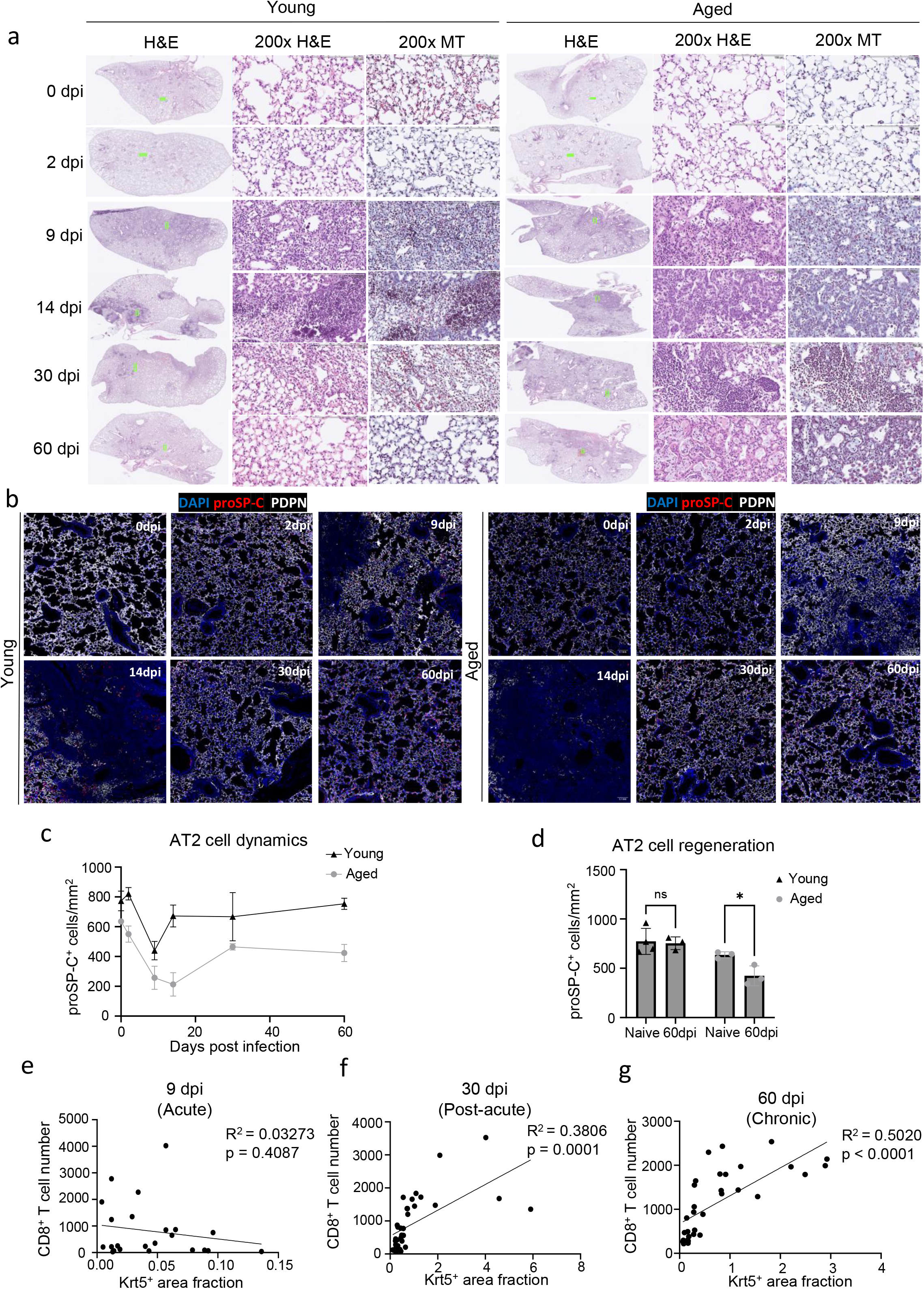
Chronic pathology and tissue sequelae after influenza infection mimics features of human PASC-PF. **(a)** Representative H&E and MT images of young and aged C57BL/6 influenza-infected mouse lungs over the course of infection. **(b)** Representative immunofluorescence images of young and aged C57BL/6 influenza-infected mouse lungs, staining alveolar epithelial cell markers (AT1-PDPN; AT2 – proSP-C) over the course of influenza infection. **(c)** Quantification of AT2 (proSP-C^+^) cell numbers in young and aged mice over the course of influenza infection (n= 3-4 per timepoint). **(d)** Quantification of AT2 (proSP-C+) cells in naïve and influenza-infected (60dpi) young and aged mouse lungs (n= 3-4 per timepoint). **(e)** Simple linear regression of CD8^+^ T cell number and Krt5^+^ area fraction in aged influenza-infected mouse lungs at 9 dpi, **(f)** 30dpi and **(g)** 60dpi. Data are expressed as mean ± SEM. Statistical analyses were conducted using a two-way ANOVA. *p < 0.05.

**Extended data Fig.7.**
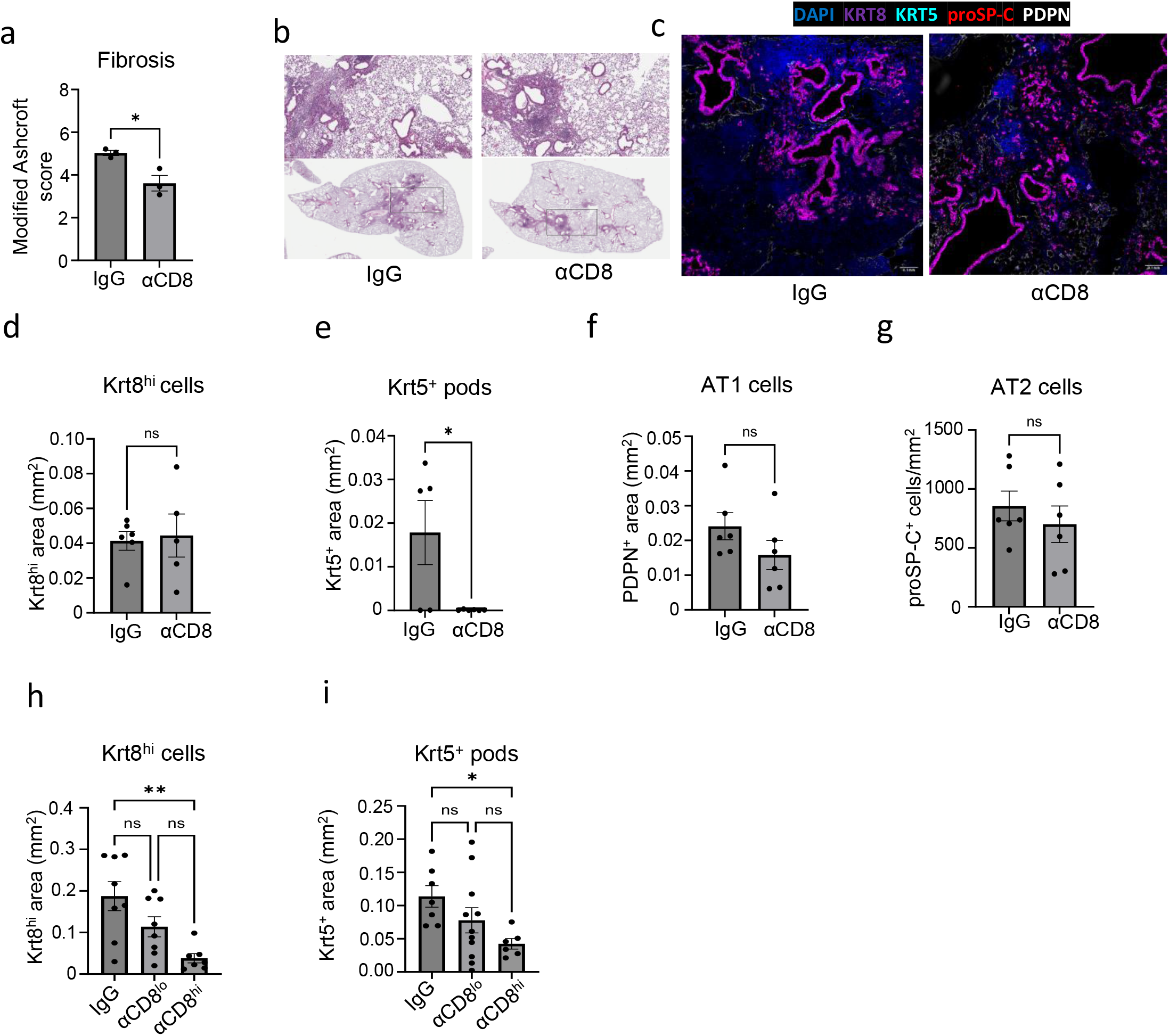
Depletion of lung-resident CD8^+^ T cells improves fibrotic disease in aged but not young mice. **(a)** Evaluation of fibrotic disease in aged influenza-infected C57BL/6 mice (60dpi) following treatment with αCD8 or control IgG Ab. (n = 3 control IgG, 3 αCD8) **(b)** Representative H&E images of young C57BL/6 lungs post influenza infection (60dpi) treated with αCD8 or control IgG Ab. **(c)** Representative immunofluorescence images of young C57BL/6 lungs post influenza infection (60dpi) treated with αCD8 or control IgG Ab. **(d)** Quantification of Krt8^hi^, **(e)** Krt5^+^, **(f)** AT1, and **(g)** AT2 cells in young influenza-infected mouse lungs, treated with αCD8 Ab or control IgG Ab. (n = 5 IgG, 5 αCD8) **(h)** Quantification of Krt8^hi^ and **(k)** Krt5^+^ area in aged influenza-infected mouse lungs (60dpi), treated with control IgG Ab, low dose αCD8 or high dose αCD8 (n = 6-8 per condition). Data are expressed as mean ± SEM. Statistical analyses were conducted using a two-tailed unpaired t-test (a,d-g), and an ordinary one-way ANOVA (h,i). *p < 0.05; **p < 0.01.

**Extended data Fig.8.**
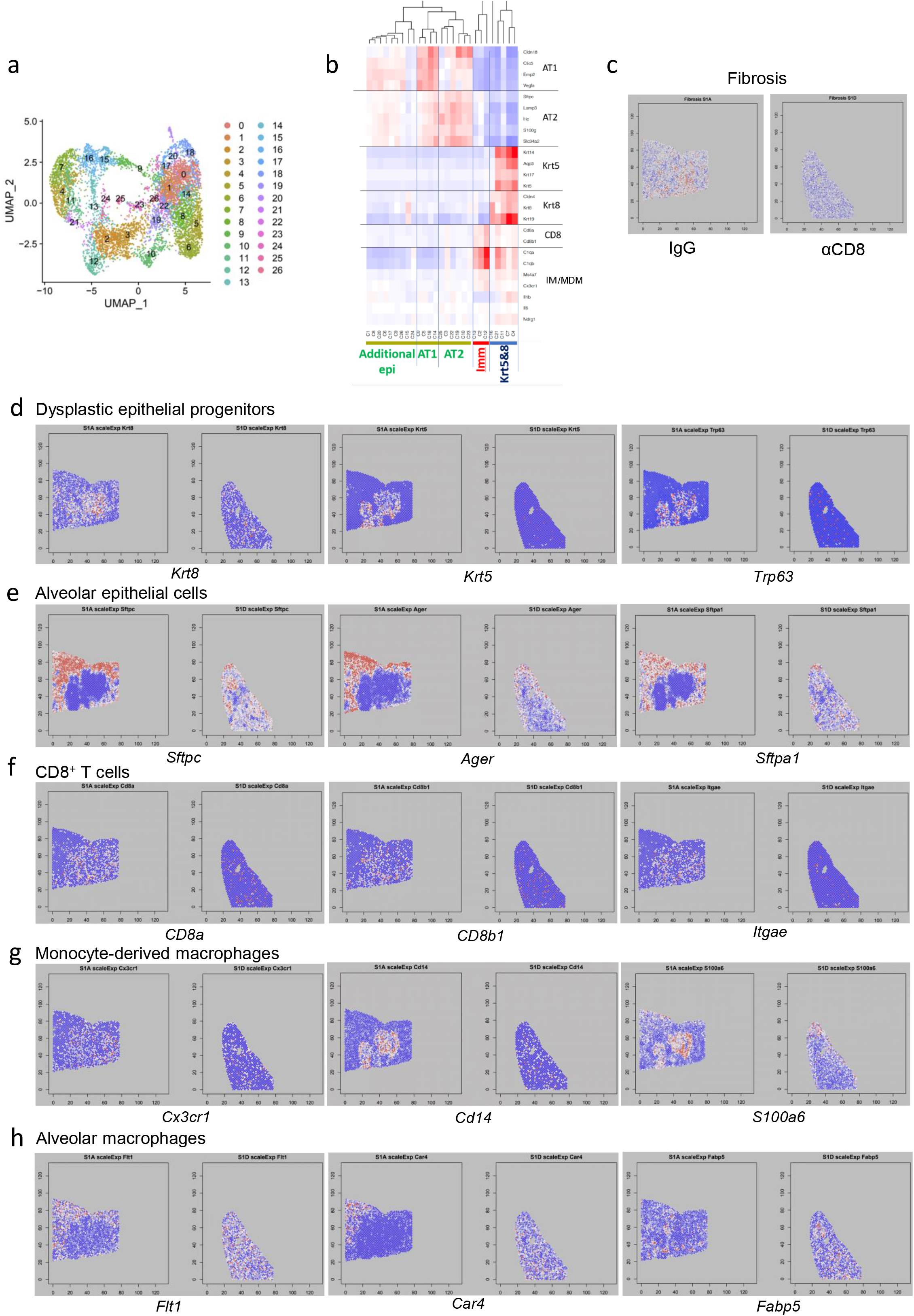
CD8^+^ T cell depletion induces widespread changes in immune and epithelial cells gene expression when evaluated by spatial transcriptomics. **(a)** UMAP visualization of spatial transcriptomics data from aged influenza-infected mice (60dpi), treated with control IgG Ab (N=2) or αCD8 (N=2). **(b)** Heatmap of gene expression across all identified clusters of spots. **(c)** Spatial map of the expression of genes associated with lung fibrosis in aged influenza-infected mice (60dpi), treated with control IgG Ab (S1A) or αCD8 (S1D). **(d)** Spatial map of the expression of dysplastic epithelial progenitors (*Krt8, Krt5, Trp63*), **(e)** alveolar epithelial (*S*ftpc1*, Ager, S*ftpa1), **(f)** CD8^+^ T cell (*CD8a, CD8b1, Itgae*), **(g)** monocyte-derived macrophages (*Cx3cr1, Cd14, S100a6*), and **(h)** alveolar macrophage (*Flt1, Car4, Fabp5*) marker genes in aged influenza-infected mice (60dpi), treated with control IgG Ab (S1A) or αCD8 (S1D).

**Extended data Fig.9.**
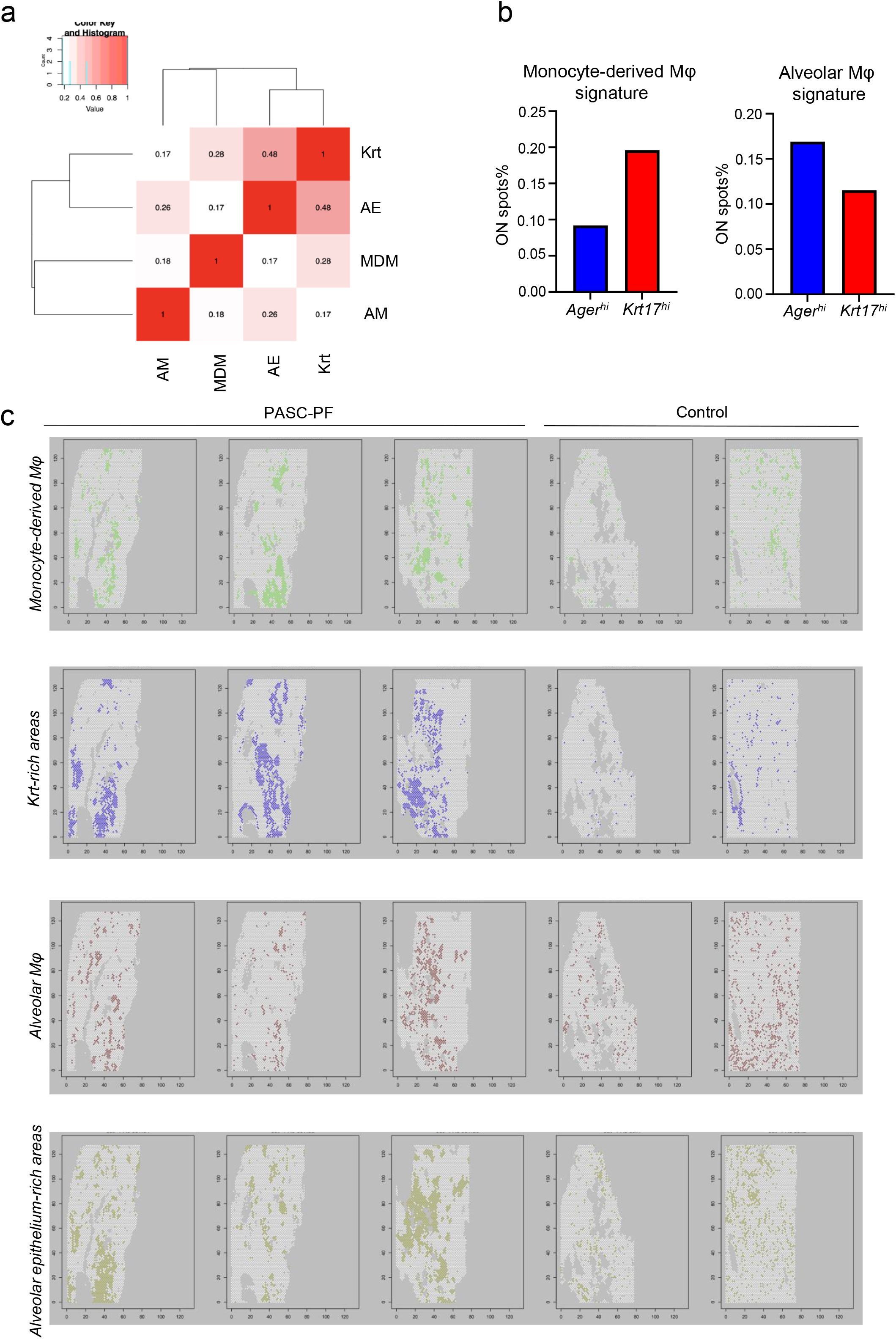
Monocyte-derived macrophages but not alveolar macrophages are enriched in areas of dysplastic repair. **(a)** Heatmap of the physical distribution of monocyte-derived macrophages (MDM), alveolar macrophages (AM), healthy alveolar epithelium (AE), and dysplastic areas (Krt) within human PASC-PF lungs. **(b)** Quantification of the proportion of spots expressing gene signatures characterizing monocyte-derived macrophages and alveolar macrophages in *Ager^hi^* and *Krt17^hi^* areas of human PASC-PF lungs. **(c)** Spatial gene expression maps of monocyte-derived macrophages, Krt-rich dysplastic areas, alveolar macrophages and healthy alveolar epithelium in human control and PASC-PF lungs.

**Extended data Fig.10.**
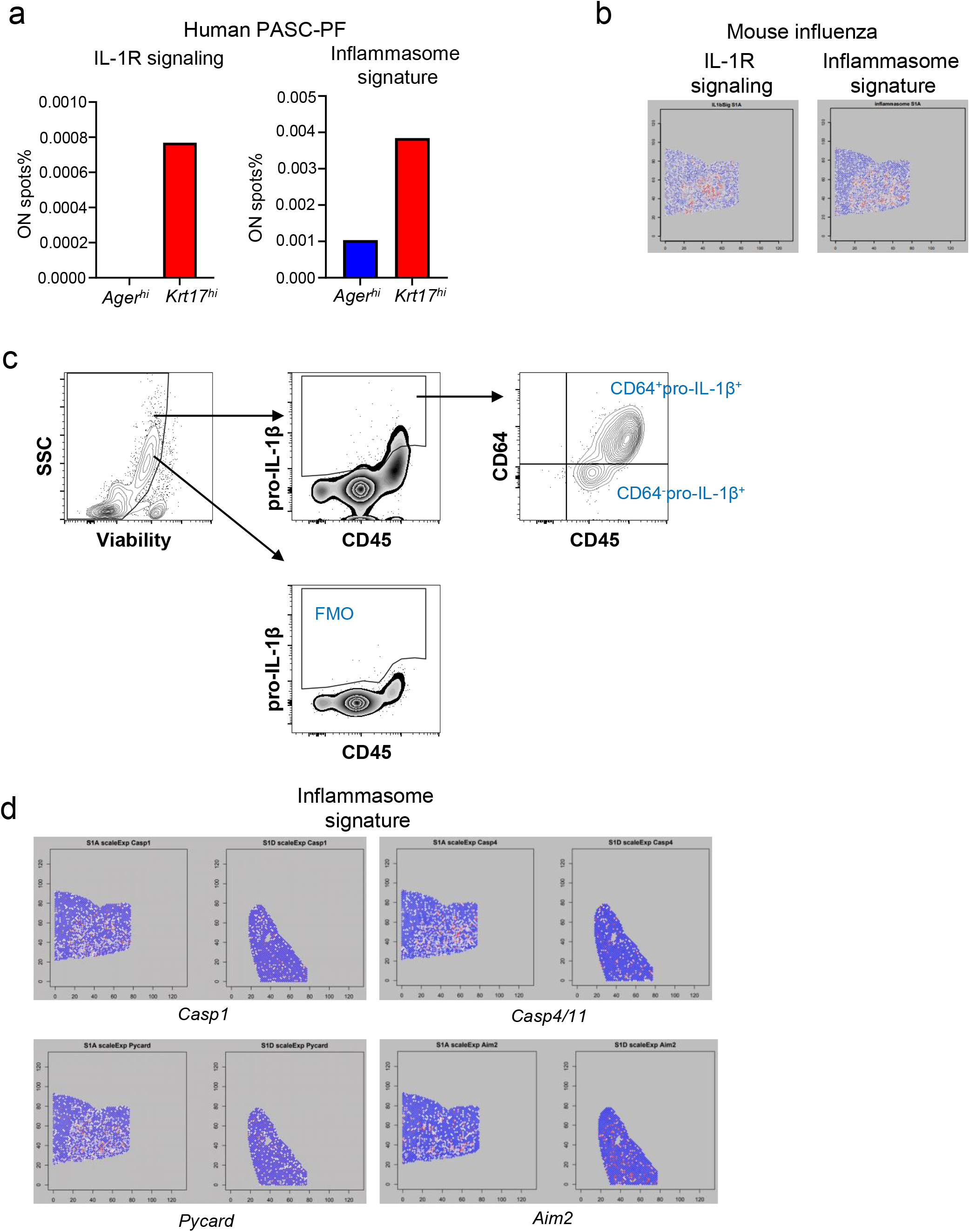
Areas of dysplastic repair are enriched with IL-1R signaling and inflammasome signatures in human PASC-PF and aged influenza-infected mouse lungs. **(a)** Quantification of the proportion of spots expressing gene signatures of IL-1R signaling and inflammasome activity in *Ager^hi^* and *Krt17^hi^* areas of human PASC-PF lungs. **(b)** Spatial gene expression maps of IL-1R signaling and inflammasome activity in aged influenza-infected mouse lungs (60dpi). **(c)** Gating strategy to identify proIL-1β^+^ cells in the lungs of influenza-infected mice by flow cytometry. **(d)** Spatial gene expression maps of inflammasome components in aged influenza-infected mice (60dpi) treated with control IgG Ab (S1A) or αCD8 (S1D).

**Extended data Fig.11.**
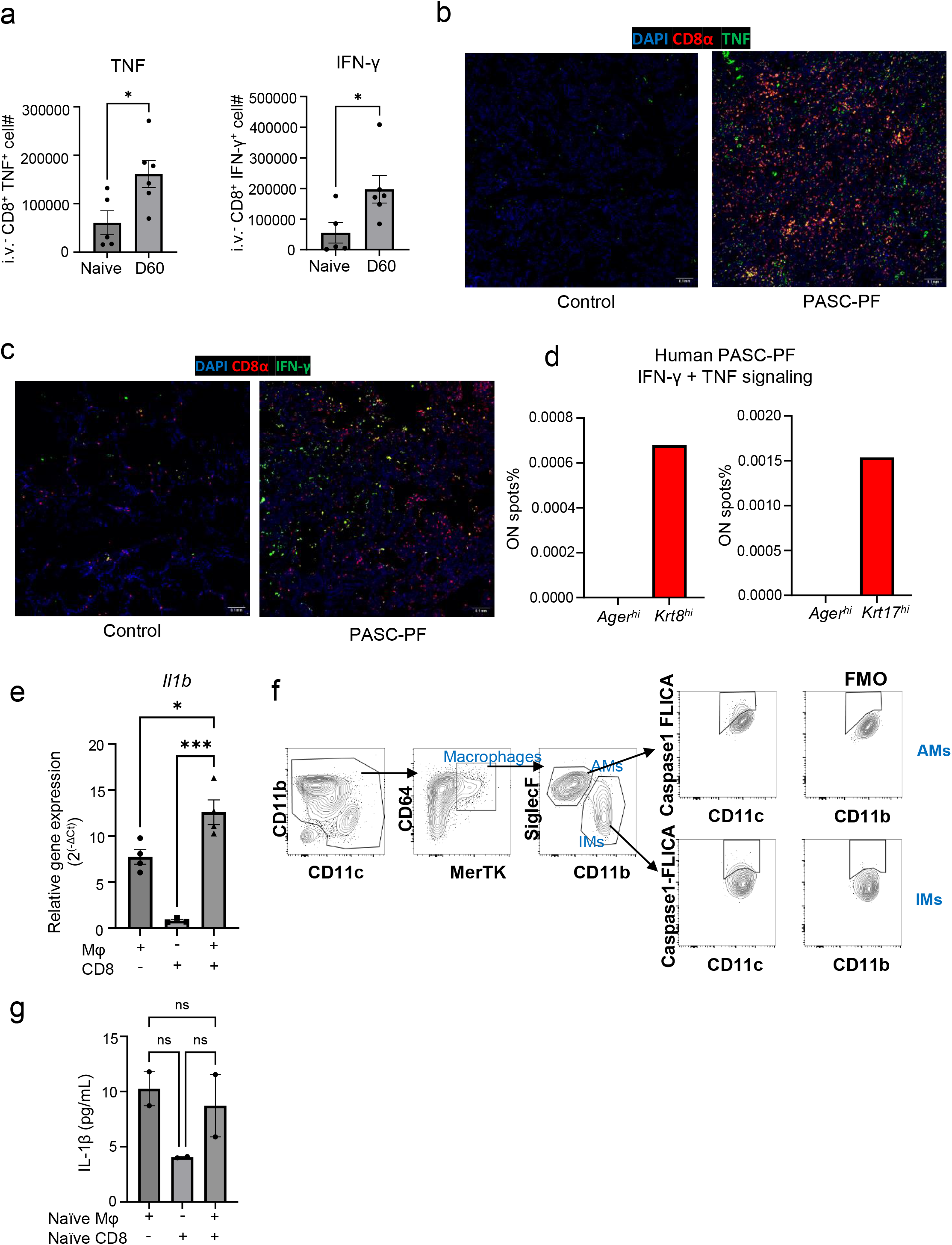
CD8^+^ T cell derived IFN-γ and TNF promotes IL-1β release. **(a)** Quantification of TNF^+^ and IFN-γ^+^ lung-resident CD8^+^ T cells in naïve and influenza-infected (60dpi) aged mouse lungs. **(b)** Representative immunofluorescence images staining for CD8^+^ T cells (CD8α) and TNF in human control and PASC-PF lungs. **(c)** Representative immunofluorescence images staining for CD8 T cells (CD8α) and IFN-γ in human control and PASC-PF lungs. **(d)** Quantification of the proportion of spots expressing gene signatures of IFN-γ + TNF signaling in healthy (*Ager^hi^*) and dysplastic areas (*Krt8^hi^* or *Krt17^+^*) within human PASC-PF lungs. **(e)** Quantification of IL-1β gene expression in following macrophage and CD8^+^ T cell coculture. **(f)** Gating strategy to identify FLICA^+^ cells following infection of aged C57BL/6 mice. **(g)** Evaluation of IL-1β release into the supernatant following isolation and coculture of macrophages and CD8^+^ T cells from naïve aged C57BL/6 mice. Data are expressed as mean ± SEM. Statistical analyses were conducted using a two-tailed unpaired t-test (a), and an ordinary one-way ANOVA (e,g). *p < 0.05; *** p <0.001.

**Extended data Fig.12.**
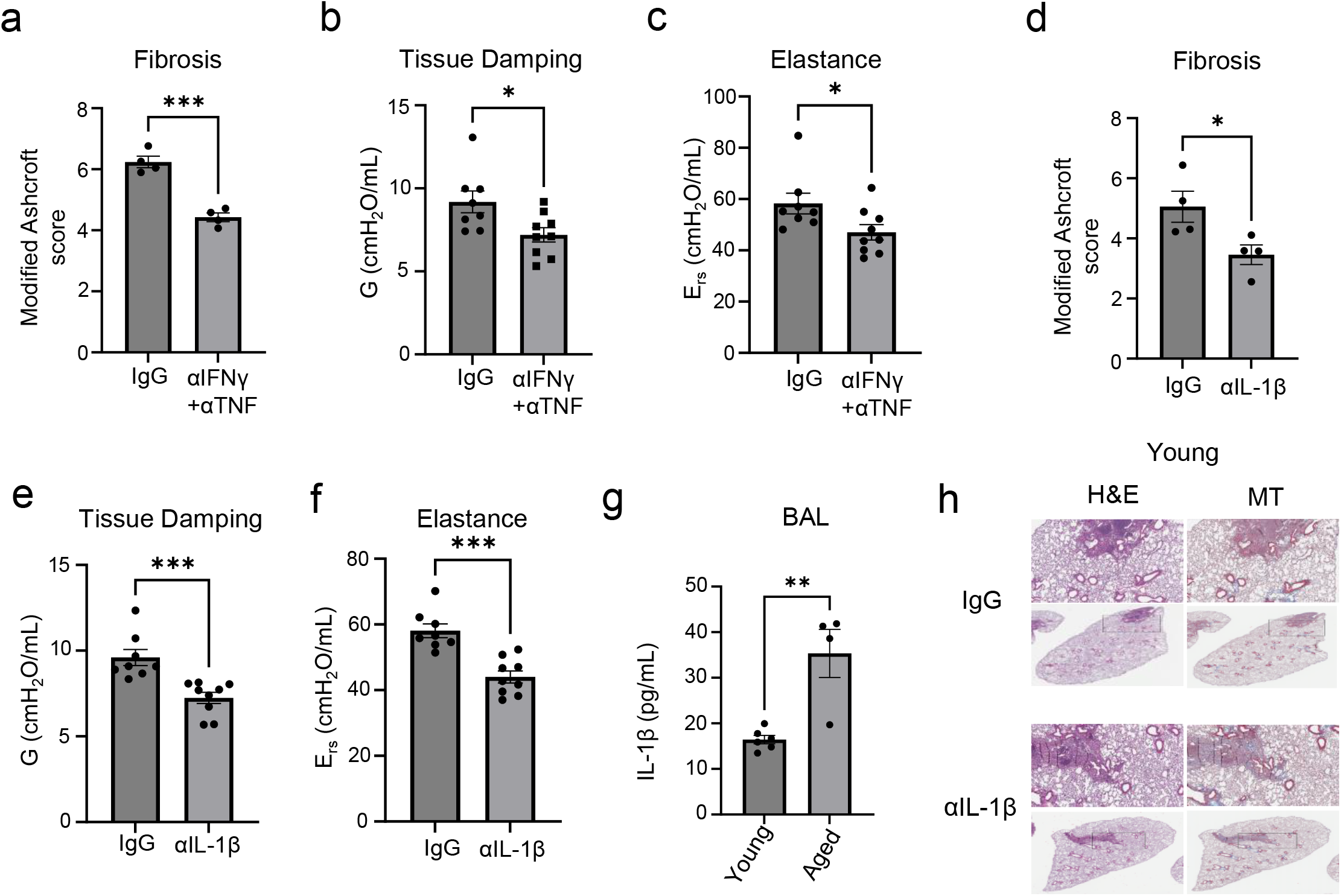
Neutralization of IFN-γ and TNF or IL-1β activity in the post-acute phase of infection improves outcomes in aged influenza-infected mice. **(a)** Evaluation of fibrotic disease in aged influenza infected C57BL/6 mice (42dpi) treated with control IgG Ab or αIFN-γ + αTNF neutralizing Ab. **(b)** Quantification of tissue damping (G) and **(c)** elastance of the respiratory system (E_rs_) in aged influenza infected C57BL/6 mice (42dpi) treated with control IgG Ab or αIFN-γ + αTNF neutralizing Ab. **(d)** Evaluation of fibrotic disease in aged influenza infected C57BL/6 mice (42dpi) treated with control IgG Ab or αIL-1β neutralizing Ab. **(e)** Quantification of tissue damping (G) and **(f)** elastance of the respiratory system (E_rs_) in aged influenza infected C57BL/6 mice (42dpi) treated with control IgG Ab or αIL-1β neutralizing Ab. **(g)** Evaluation of IL-1β levels in the BAL fluid of young and aged influenza-infected mice (42dpi). **(h)** Representative H&E and MT images of young C57BL/6 lungs post influenza infection (42pi) treated with αIL-1β neutralizing Ab or control IgG Ab. Data are expressed as mean ± SEM. Statistical analyses were conducted using a two-tailed unpaired t-test. *p < 0.05; **p < 0.01; ***p < 0.001.

**Extended data Fig.13.**
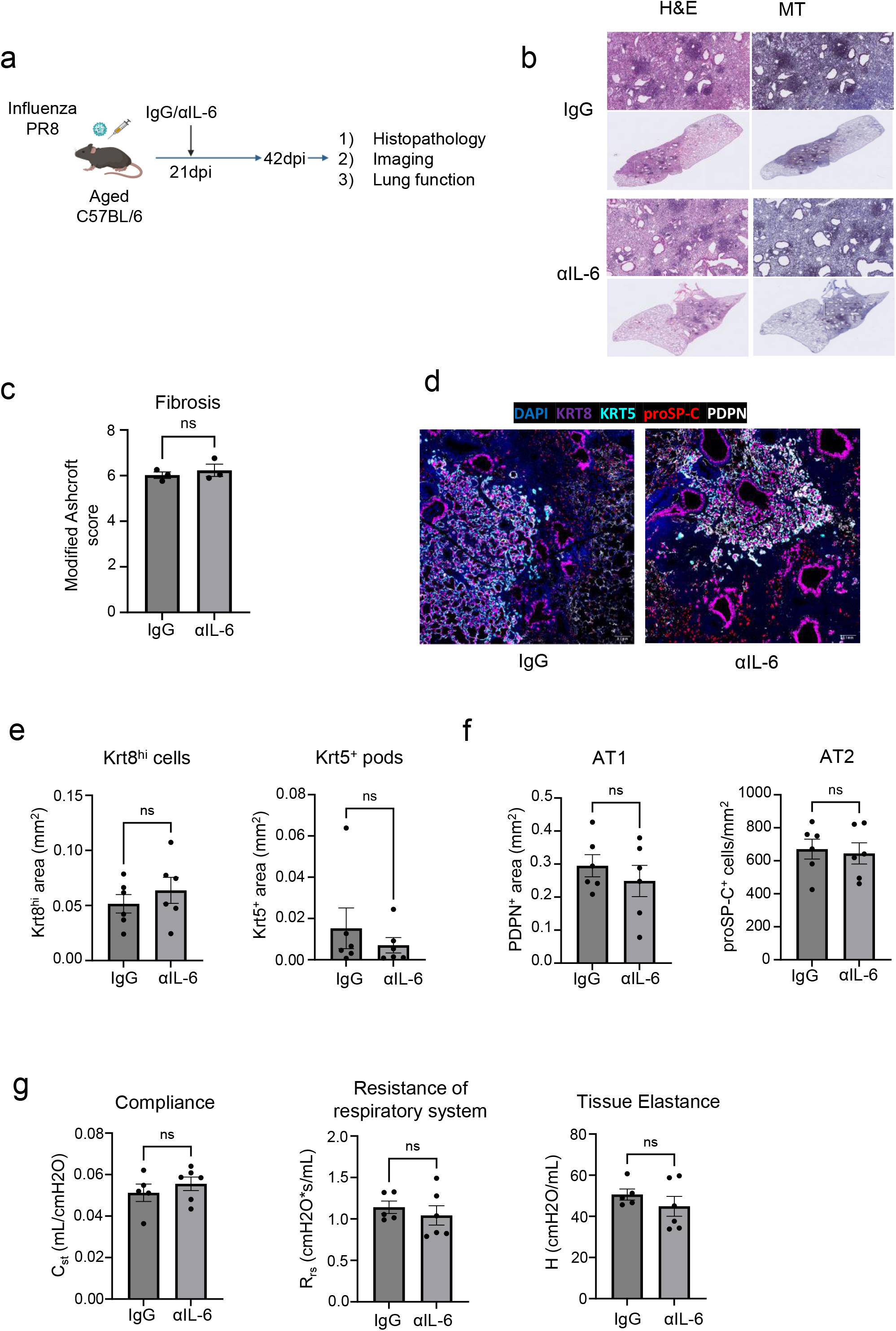
Neutralization of IL-6 activity in the post-acute phase of influenza infection does not improve long-term outcomes. **(a)** Experimental design for *in vivo* IL-6 neutralization post PR8 influenza infection. **(b)** Representative H&E and MT images of aged C57BL/6 lungs post PR8 influenza infection (42dpi) treated with αIL-6 neutralizing Ab or control IgG Ab. **(c).** Evaluation of fibrotic disease in aged influenza-infected (42dpi) mice treated with αIL-6 neutralizing Ab or control IgG Ab. **(d)** Representative immunofluorescence images staining AT1 (PDPN^+^), AT2 (proSP-C^+^) and epithelial progenitors (Krt5^+^ and Krt8^hi^) in aged influenza-infected mice (42dpi) treated with αIL-6 neutralizing Ab or control IgG Ab. **(e)** Quantification of Krt8^hi^ and Krt5^+^ area and **(f)** AT1 (PDPN^+^) and AT2 (proSP-C^+^) cells in aged influenza-infected mice treated with αIL-6 neutralizing Ab or control IgG Ab (n= 6 control IgG, 6 αIL-6). **(g)** Evaluation of static compliance (C_st_), resistance of the respiratory system (R_rs_), and tissue elastance (H) in aged influenza-infected mice (42dpi) treated with αIL-6 neutralizing Ab or control IgG Ab (n= 5 control IgG, 6 αIL-6). Data are expressed as mean ± SEM. Statistical analyses were conducted using a two-tailed unpaired t-test. *p < 0.05.

**Extended data Fig.14.**
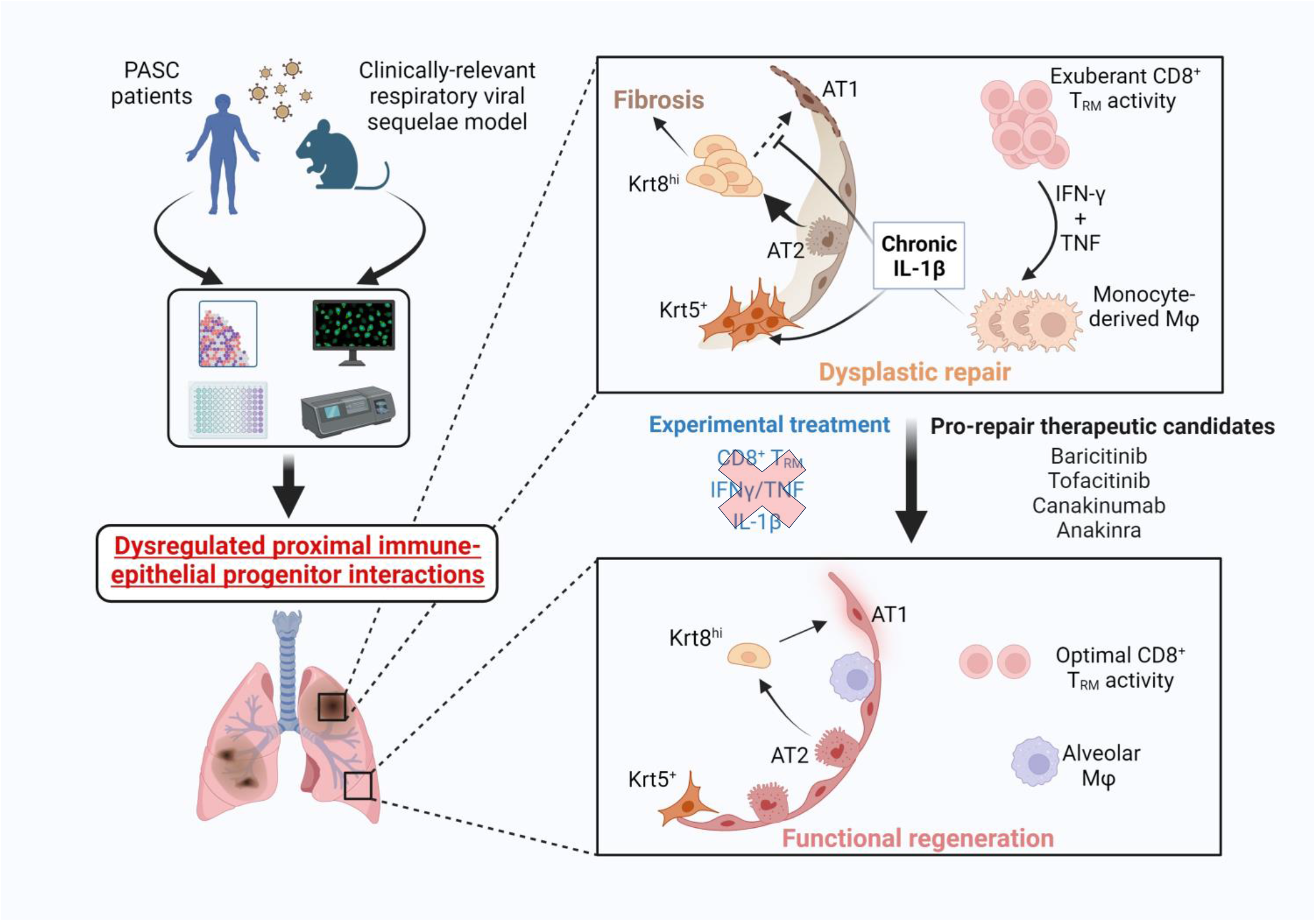
Dysregulated immune-epithelial progenitor interactions drive post-viral sequelae in PASC.

**Supplementary Table1.**
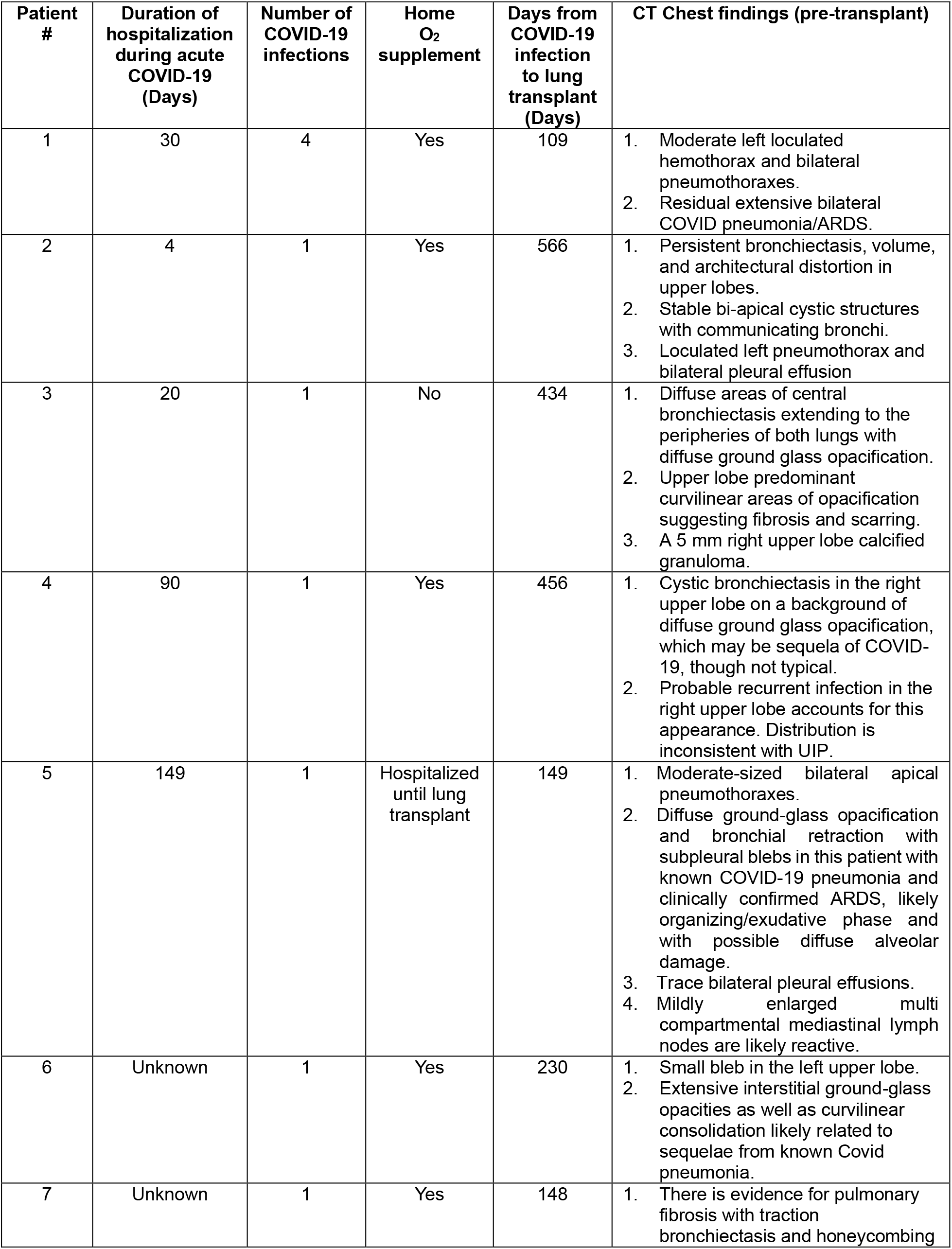
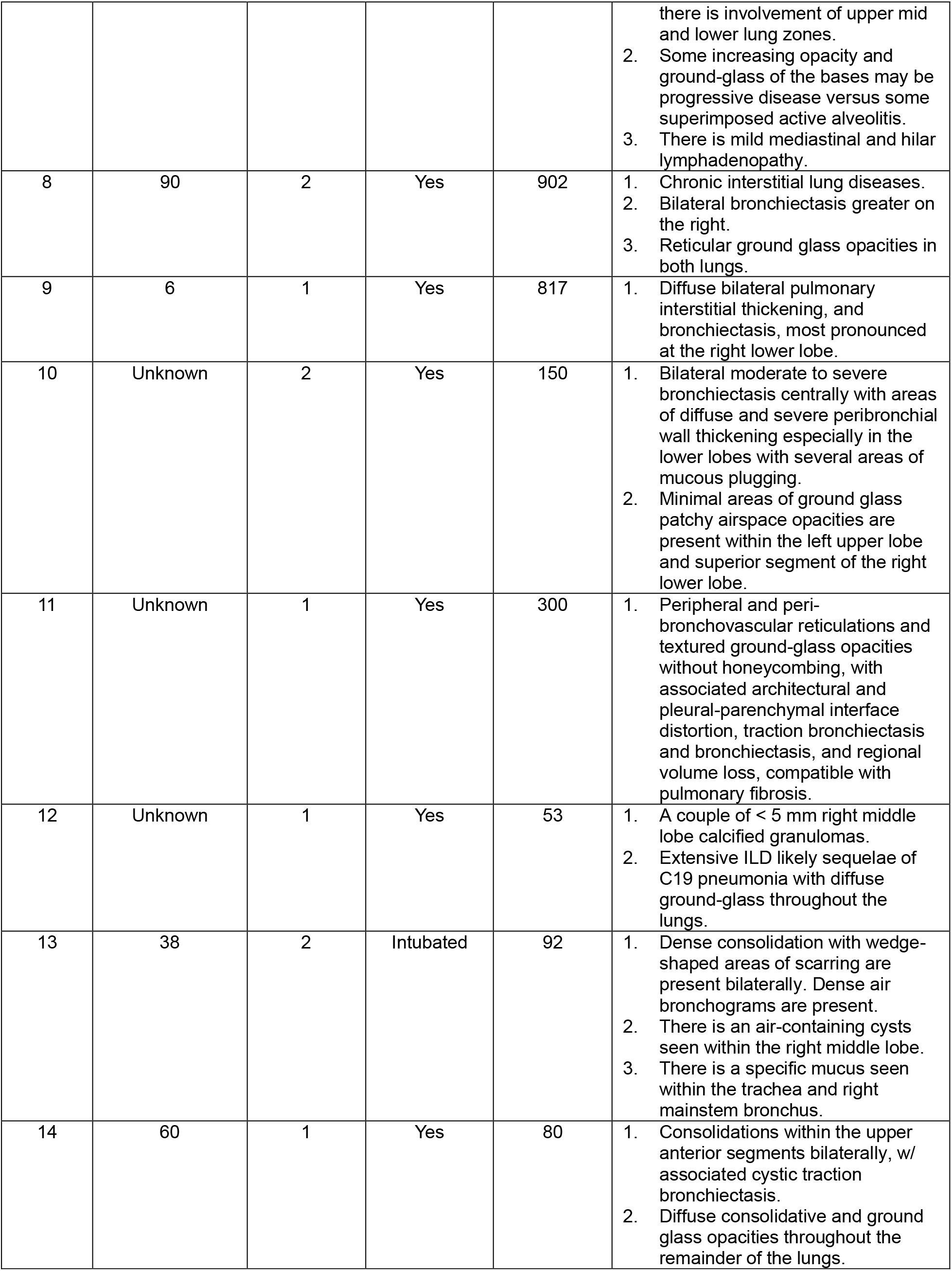
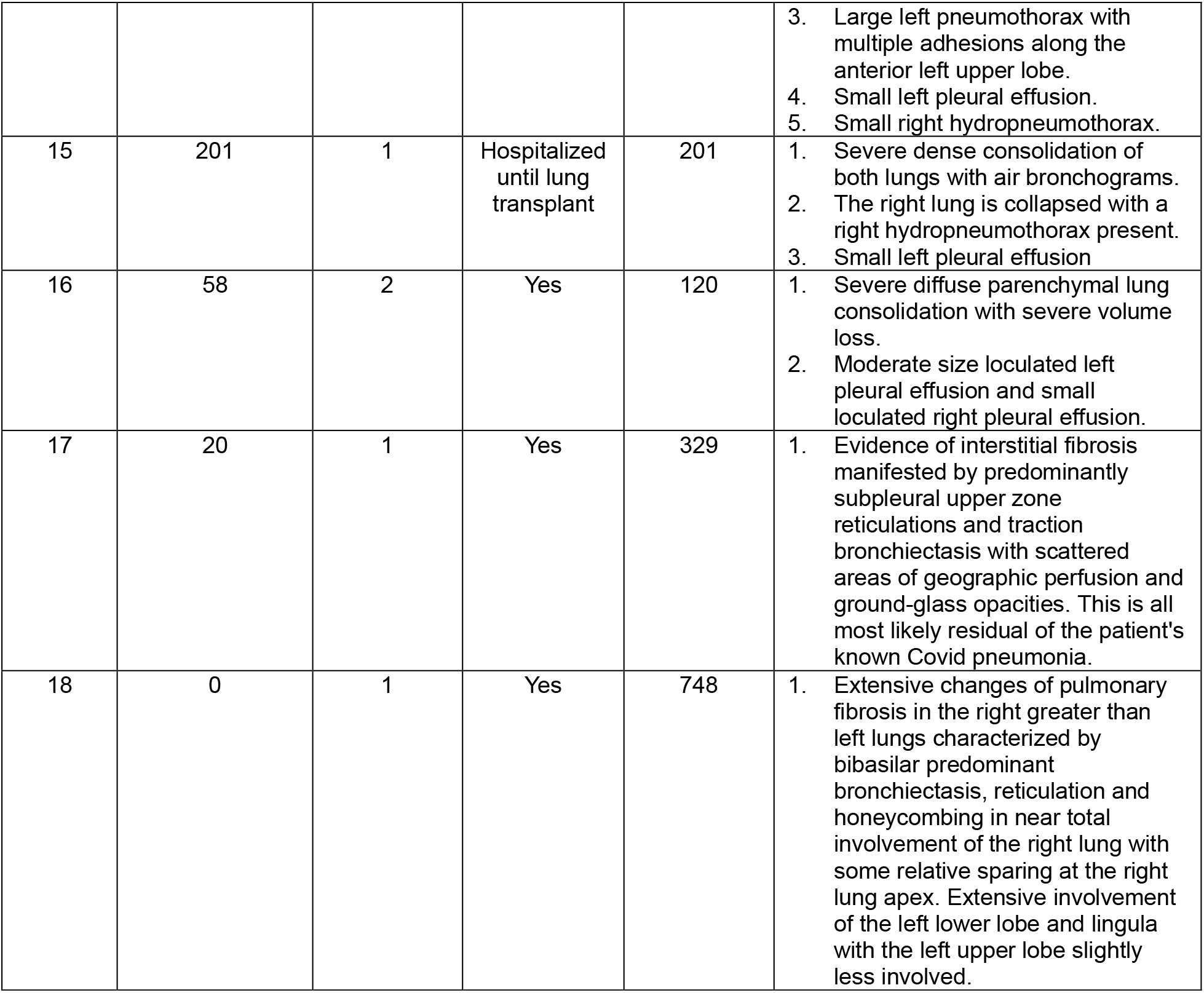
Clinical information of the PASC-PF cohort.

| Patient # | Duration of hospitalization during acute COVID-19 (Days) | Number of COVID-19 infections | Home O<sub>2</sub> supplement | Days from COVID-19 infection to lung transplant (Days) | CT Chest findings (pre-transplant) |
| --- | --- | --- | --- | --- | --- |
| 1 | 30 | 4 | Yes | 109 | 1. Moderate left loculated hemothorax and bilateral pneumothoraxes.<br>2. Residual extensive bilateral COVID pneumonia/ARDS. |
| 2 | 4 | 1 | Yes | 566 | 1. Persistent bronchiectasis, volume, and architectural distortion in upper lobes.<br>2. Stable bi-apical cystic structures with communicating bronchi.<br>3. Loculated left pneumothorax and bilateral pleural effusion |
| 3 | 20 | 1 | No | 434 | 1. Diffuse areas of central bronchiectasis extending to the peripheries of both lungs with diffuse ground glass opacification.<br>2. Upper lobe predominant curvilinear areas of opacification suggesting fibrosis and scarring.<br>3. A 5 mm right upper lobe calcified granuloma. |
| 4 | 90 | 1 | Yes | 456 | 1. Cystic bronchiectasis in the right upper lobe on a background of diffuse ground glass opacification, which may be sequela of COVID-19, though not typical.<br>2. Probable recurrent infection in the right upper lobe accounts for this appearance. Distribution is inconsistent with UIP. |
| 5 | 149 | 1 | Hospitalized until lung transplant | 149 | 1. Moderate-sized bilateral apical pneumothoraxes.<br>2. Diffuse ground-glass opacification and bronchial retraction with subpleural blebs in this patient with known COVID-19 pneumonia and clinically confirmed ARDS, likely organizing/exudative phase and with possible diffuse alveolar damage.<br>3. Trace bilateral pleural effusions.<br>4. Mildly enlarged multi compartmental mediastinal lymph nodes are likely reactive. |
| 6 | Unknown | 1 | Yes | 230 | 1. Small bleb in the left upper lobe.<br>2. Extensive interstitial ground-glass opacities as well as curvilinear consolidation likely related to sequelae from known Covid pneumonia. |
| 7 | Unknown | 1 | Yes | 148 | 1. There is evidence for pulmonary fibrosis with traction bronchiectasis and honeycombing<br>there is involvement of upper mid and lower lung zones.<br>2. Some increasing opacity and ground-glass of the bases may be progressive disease versus some superimposed active alveolitis.<br>3. There is mild mediastinal and hilar lymphadenopathy. |
| 8 | 90 | 2 | Yes | 902 | 1. Chronic interstitial lung diseases.<br>2. Bilateral bronchiectasis greater on the right.<br>3. Reticular ground glass opacities in both lungs. |
| 9 | 6 | 1 | Yes | 817 | 1. Diffuse bilateral pulmonary interstitial thickening, and bronchiectasis, most pronounced at the right lower lobe. |
| 10 | Unknown | 2 | Yes | 150 | 1. Bilateral moderate to severe bronchiectasis centrally with areas of diffuse and severe peribronchial wall thickening especially in the lower lobes with several areas of mucous plugging.<br>2. Minimal areas of ground glass patchy airspace opacities are present within the left upper lobe and superior segment of the right lower lobe. |
| 11 | Unknown | 1 | Yes | 300 | 1. Peripheral and peri-bronchovascular reticulations and textured ground-glass opacities without honeycombing, with associated architectural and pleural-parenchymal interface distortion, traction bronchiectasis and bronchiectasis, and regional volume loss, compatible with pulmonary fibrosis. |
| 12 | Unknown | 1 | Yes | 53 | 1. A couple of < 5 mm right middle lobe calcified granulomas.<br>2. Extensive ILD likely sequelae of C19 pneumonia with diffuse ground-glass throughout the lungs. |
| 13 | 38 | 2 | Intubated | 92 | 1. Dense consolidation with wedge-shaped areas of scarring are present bilaterally. Dense air bronchograms are present.<br>2. There is an air-containing cysts seen within the right middle lobe.<br>3. There is a specific mucus seen within the trachea and right mainstem bronchus. |
| 14 | 60 | 1 | Yes | 80 | 1. Consolidations within the upper anterior segments bilaterally, w/ associated cystic traction bronchiectasis.<br>2. Diffuse consolidative and ground glass opacities throughout the remainder of the lungs. |
|  |  |  |  |  | 3. Large left pneumothorax with multiple adhesions along the anterior left upper lobe.<br>4. Small left pleural effusion.<br>5. Small right hydropneumothorax. |
| 15 | 201 | 1 | Hospitalized until lung transplant | 201 | 1. Severe dense consolidation of both lungs with air bronchograms.<br>2. The right lung is collapsed with a right hydropneumothorax present.<br>3. Small left pleural effusion |
| 16 | 58 | 2 | Yes | 120 | 1. Severe diffuse parenchymal lung consolidation with severe volume loss.<br>2. Moderate size loculated left pleural effusion and small loculated right pleural effusion. |
| 17 | 20 | 1 | Yes | 329 | 1. Evidence of interstitial fibrosis manifested by predominantly subpleural upper zone reticulations and traction bronchiectasis with scattered areas of geographic perfusion and ground-glass opacities. This is all most likely residual of the patient's known Covid pneumonia. |
| 18 | 0 | 1 | Yes | 748 | 1. Extensive changes of pulmonary fibrosis in the right greater than left lungs characterized by bibasilar predominant bronchiectasis, reticulation and honeycombing in near total involvement of the right lung with some relative sparing at the right lung apex. Extensive involvement of the left lower lobe and lingula with the left upper lobe slightly less involved. |

**Supplementary Table 2.** Clinical Information of COVID-19 Convalescent Subjects.

|  | **Normal PFT<sup>1</sup>**<br>N = 29 | **Abnormal PFT<sup>1</sup>**<br>N = 39 | **p-value<sup>2</sup>** |
| --- | --- | --- | --- |
| **Age (years)** | 51 (15) | 50 (12) | 0.12 |
| **Sex** |  |  | 0.80 |
| Female | 13 (45%) | 16 (41%) |  |
| Male | 16 (55%) | 23 (59%) |  |
| **Race** |  |  | 0.60 |
| White | 15 (52%) | 23 (59%) |  |
| Black | 5 (17%) | 8 (21%) |  |
| Other | 9 (31%) | 8 (21%) |  |
| **Ethnicity** |  |  | 0.60 |
| Hispanic | 8 (28%) | 13 (33%) |  |
| Non-Hispanic | 21 (72%) | 26 (67%) |  |
| **Admitted to Hospital** | 22 (76%) | 38 (97%) | 0.009 |
| Duration (Days) | 14 (16) | 21 (25) | 0.14 |
| **Time since Illness (Days)<sup>3</sup>** | 113 (110) | 98 (104) | 0.80 |

<sup>1</sup>Median (SD); n (%)
<sup>2</sup>Wilcoxon rank sum test; Pearson's Chi-squared test; Fisher's exact test
<sup>3</sup>Earliest date: First symptoms, positive test, hospital admission

**Supplementary Table 3.**
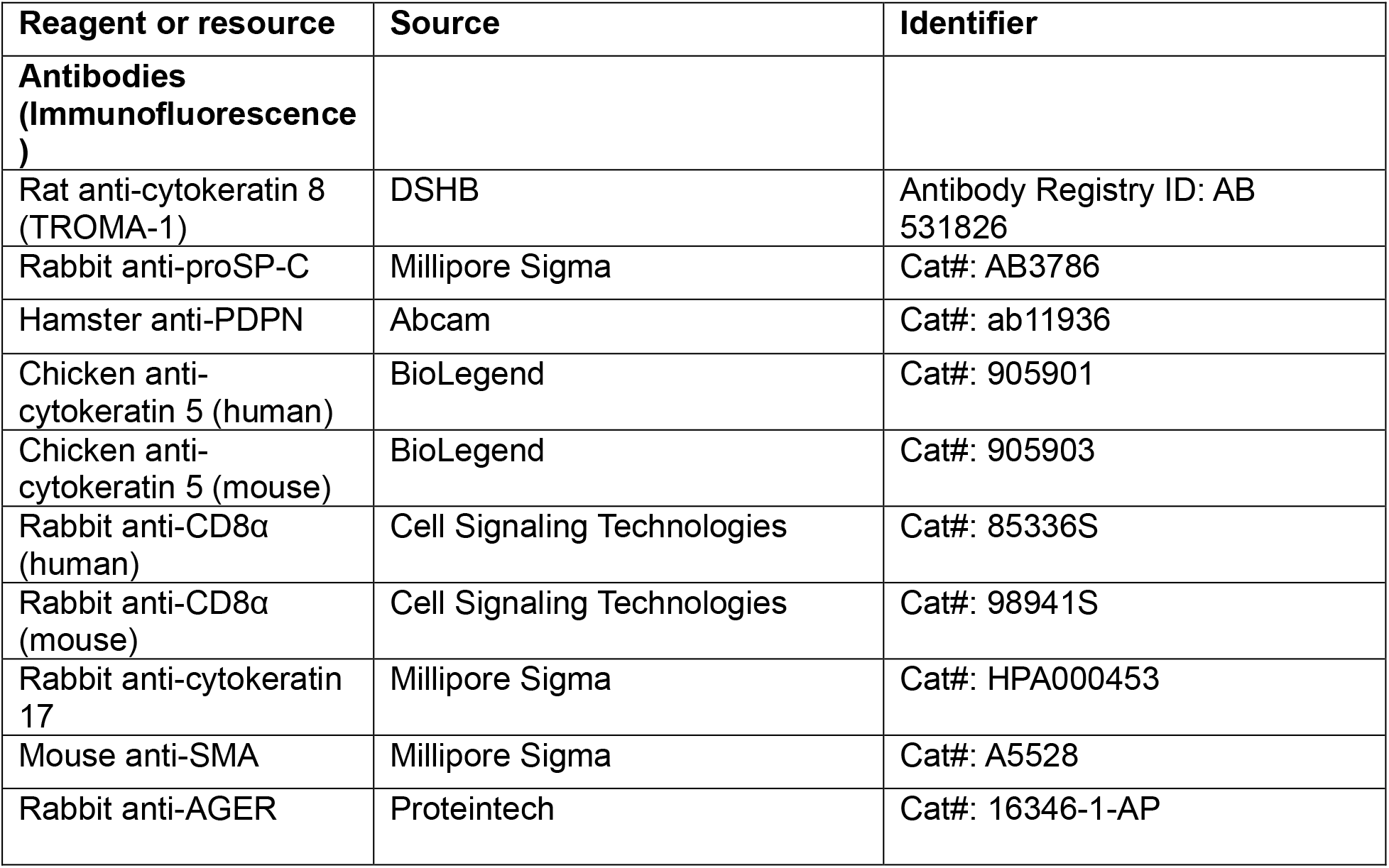
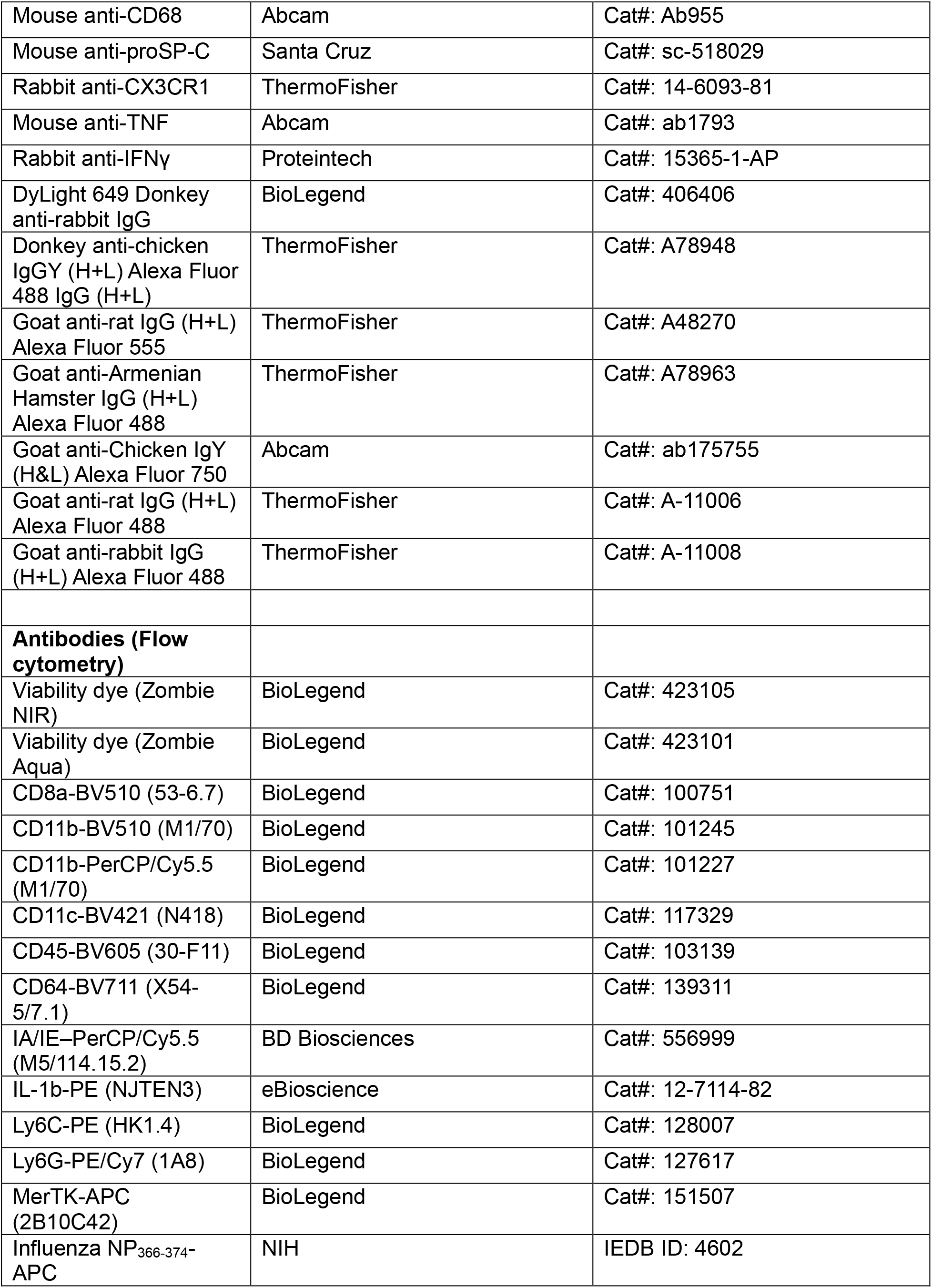
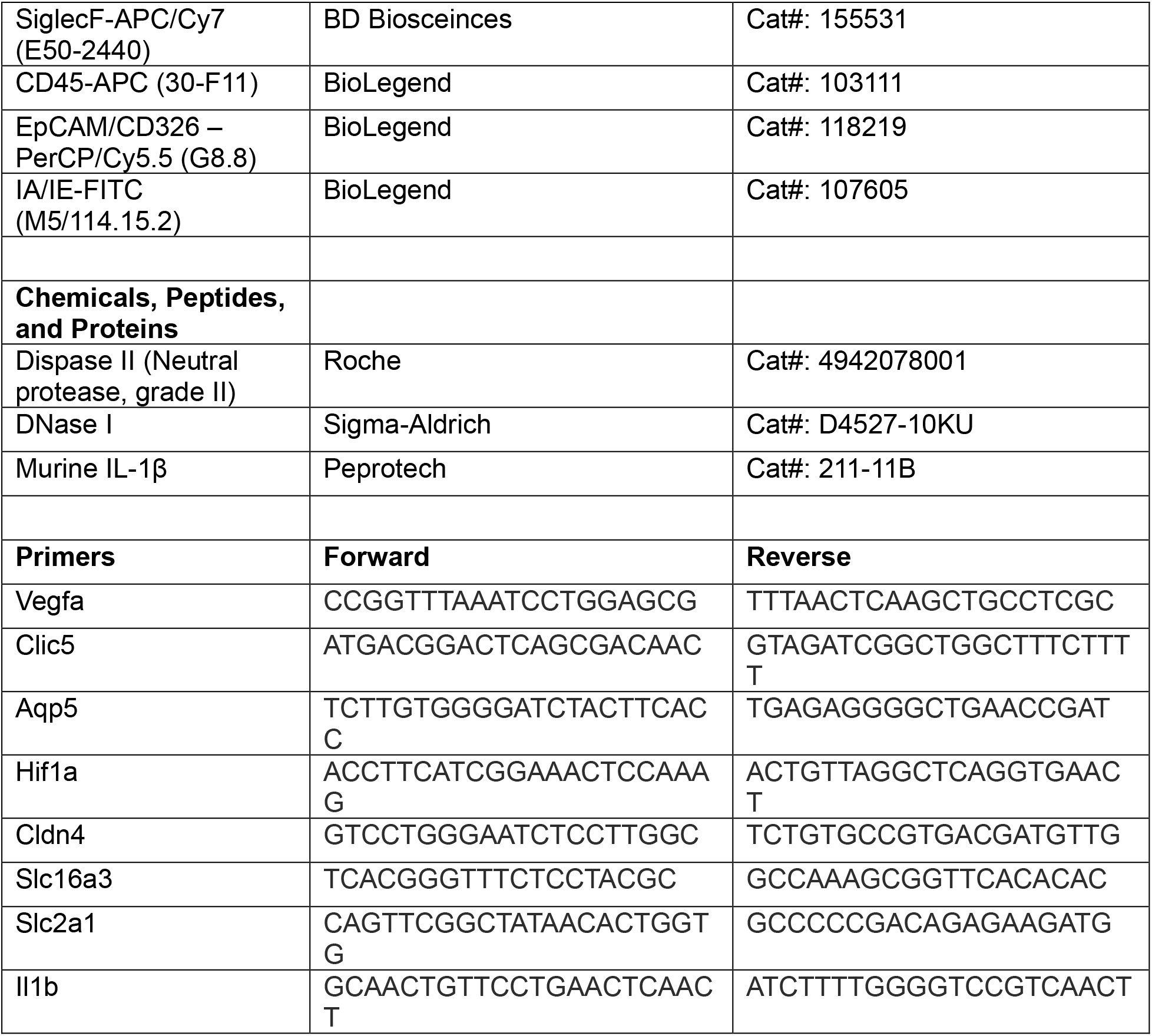
Key resources.

**Supplementary Table 4.** Spatial transcriptomics gene sets.

| Cell type/Signaling pathway | Marker genes/source |
| --- | --- |
| **MOUSE** |  |
| CD8 T cells | *CD8a*, *CD8b1*, *Itgae*, *Cd3d*, *Trbc1*, *Ccl5*, *Gzma*, *Gzmb* |
| Monocyte derived macrophages | *Cd14*, *S100a6*, *Apoe*, *Mafb*, *Vcan*, *Fn1*, *Stat1* |
| Alveolar macrophages | *Sparc*, *Flt1*, *Fabp5*, *Car4*, *Krt79* |
| Alveolar epithelium | *Sftpc*, *Clic5*, *Emp2*, *Hopx*, *Spock2*, *Sftpa1*, *Cldn18* |
| Krt-rich dysplastic repair | *Krt5*, *Krt8*, *Krt17*, *Trp53*, *Cldn4* |
| ADI/PATS/DATP signature | (20), (21), (22) |
| Aberrant basaloid signature | (23), (24) |
| Fibrosis | WP_Lung_Fibrosis |
| IL-1R signaling | BIOCARTA_IL1R_PATHWAY (GSEA; Mouse) |
| Inflammasome signature | REACTOME_INFLAMMASOMES (GSEA; Mouse) |
| **HUMAN** |  |
| CD8 T cells | CD8A, CCL5, GZMH, GZMA, GZMK, GZMB, GNLY |
| Monocyte derived macrophages | CD14, RNASE1, S100A8, SPP1, TYMP, MS4A6A, FCGR2B |
| Alveolar macrophages | GPD1, INHBA, MME, APOC1, CD52, PCOLCE2, PPARG, FABP4, SCD |
| Alveolar epithelium | AGER, CLDN18, ABCA3, LAMP3, SFTPA1, SFTPD, SFTPC, CLIC5, AQP4 |
| Krt-rich dysplastic repair | KRT8, KRT5, KRT17, MMP7, ITGB6, AQP3, KRT19, MMP1, S100A2 |
| IL-1R signaling | BIOCARTA_IL1R_PATHWAY (GSEA; Human) |
| Inflammasome signature | REACTOME_INFLAMMASOMES (GSEA; Human) |
| IFN + TNF signaling | REACTOME_INTERFERON_GAMMA_SIGNALLING (GSEA Human),<br>HALLMARK_TNFA_SIGNALING_VIA_NFKB (GSEA; Human) |

